# Whole genome sequencing of pharmacogenetic drug response in racially and ethnically diverse children with asthma

**DOI:** 10.1101/128116

**Authors:** Angel C.Y. Mak, Marquitta J. White, Zachary A. Szpiech, Walter L. Eckalbar, Sam S. Oh, Maria Pino-Yanes, Donglei Hu, Pagé Goddard, Scott Huntsman, Joshua Galanter, Dara G. Torgerson, Ann Chen Wu, Blanca E. Himes, Soren Germer, Julia M. Vogel, Karen L. Bunting, Celeste Eng, Sandra Salazar, Kevin L. Keys, Jennifer Liberto, Thomas J. Nuckton, Thomas A. Nguyen, Pui-Yan Kwok, Albert M. Levin, Juan C. Celedón, Erick Forno, Hakon Hakonarson, Patrick M. Sleiman, Amber Dahlin, Kelan G. Tantisira, Scott T. Weiss, Denise Serebrisky, Emerita Brigino-Buenaventura, Harold J. Farber, Kelley Meade, Michael A. Lenoir, Pedro C. Avila, Saunak Sen, Shannon M. Thyne, William Rodriguez-Cintron, Cheryl A. Winkler, Andrés Moreno-Estrada, Karla Sandoval, Jose R. Rodriguez-Santana, Rajesh Kumar, L. Keoki Williams, Nadav Ahituv, Elad Ziv, Max A. Seibold, Robert B. Darnell, Noah Zaitlen, Ryan D. Hernandez, Esteban G. Burchard on behalf of the Trans-Omics for Precision Medicine Whole Genome Sequencing Program (TOPMed)

**Affiliations:** Department of Medicine, University of California San Francisco, San Francisco, California, USA.; Department of Bioengineering and Therapeutic Sciences, University of California San Francisco, San Francisco, California, USA.; Research Unit, Hospital Universitario N.S. de Candelaria, Universidad de La Laguna, Santa Cruz de Tenerife, Spain.; CIBER de Enfermedades Respiratorias, Instituto de Salud Carlos III, Madrid, Spain.; Channing Division of Network Medicine, Department of Medicine, Brigham and Women's Hospital and Harvard Medical School, Boston, Massachusetts, USA.; Precision Medicine Translational Research (PRoMoTeR) Center, Department of Population Medicine, Harvard Medical School and Pilgrim Health Care Institute, Boston, Massachusetts, USA.; Department of Biostatistics, Epidemiology and Informatics, University of Pennsylvania, Philadelphia, Pennsylvania, USA.; New York Genome Center, New York, New York, USA.; Cardiovascular Research Institute, University of California San Francisco, San Francisco, California, USA.; Institute for Human Genetics, University of California San Francisco, San Francisco, California, USA.; Department of Public Health Sciences, Henry Ford Health System, Detroit Michigan, USA.; Division of Pediatric Pulmonary Medicine, Allergy and Immunology, University of Pittsburgh School of Medicine, Pittsburgh, Pennsylvania, USA.; Center for Applied Genomics, The Children’s Hospital of Philadelphia Research Institute, Philadelphia, Pennsylvania, USA.; Department of Pediatrics, Perelman School of Medicine, University of Pennsylvania, Philadelphia, Pennsylvania, USA.; Pediatric Pulmonary Division, Jacobi Medical Center, Bronx, New York, USA.; Department of Allergy and Immunology, Kaiser Permanente Vallejo Medical Center, Vallejo, California, USA.; Department of Pediatrics, Baylor College of Medicine and Texas Children’s Hospital, Houston, Texas, USA.; Children’s Hospital and Research Center, Oakland, California, USA.; Bay Area Pediatrics, Oakland, California, USA.; Department of Medicine, Northwestern University, Chicago, Illinois, USA.; Department of Pediatrics, David Geffen School of Medicine, University of California, Los Angeles, Los Angeles, California, USA.; Veterans Caribbean Health Care System, San Juan, Puerto Rico.; Basic Science Laboratory, Center for Cancer Research, National Cancer Institute, Leidos Biomedical Research, Frederick National Laboratory, Frederick, Maryland, USA.; National Laboratory of Genomics for Biodiversity (UGA-LANGEBIO), CINVESTAV, Irapuato, Guanajuato, Mexico.; Centro de Neumologia Pediatrica, San Juan, Puerto Rico.; Feinberg School of Medicine’s Division of Allergy and Immunology, Northwestern University, Chicago, Illinois, USA.; Ann & Robert H. Lurie Children’s Hospital of Chicago, Chicago, Illinois, USA.; Department of Internal Medicine, Henry Ford Health System, Detroit, Michigan, USA.; Center for Health Policy and Health Services Research, Henry Ford Health System, Detroit, Michigan, USA.; Center for Genes, Environment and Health, Department of Pediatrics, National Jewish Health, Denver, Colorado, USA.; Laboratory of Molecular Neuro-Oncology, The Rockefeller University, New York, New York, USA.; Howard Hughes Medical Institute, The Rockefeller University, New York, New York, USA.; Quantitative Biosciences Institute, University of California San Francisco, San Francisco, California, USA.

## Abstract

Asthma is the most common chronic disease of children, with significant racial/ethnic differences in prevalence, morbidity, mortality and therapeutic response. Albuterol, a bronchodilator medication, is the first-line therapy for asthma treatment worldwide. We performed the largest whole genome sequencing (WGS) pharmacogenetics study to date using data from 1,441 minority children with asthma who had extremely high or low bronchodilator drug response (BDR). We identified population-specific and shared pharmacogenetic variants associated with BDR, including genome-wide significant (p < 3.53 x 10^-7^) and suggestive (p < 7.06 x 10^-6^) loci near genes previously associated with lung capacity (*DNAH5*), immunity (*NFKB1* and *PLCB1*), and β-adrenergic signaling pathways (*ADAMTS3* and *COX18*). Functional analyses centered on *NFKB1* revealed potential regulatory function of our BDR-associated SNPs in bronchial smooth muscle cells. Specifically, these variants are in linkage disequilibrium with SNPs in a functionally active enhancer, and are also expression quantitative trait loci (eQTL) for a neighboring gene, *SLC39A8*. Given the lack of other asthma study populations with WGS data on minority children, replication of our rare variant associations is infeasible. We attempted to replicate our common variant findings in five independent studies with GWAS data. The age-specific associations previously found in asthma and asthma-related traits suggest that the over-representation of adults in our replication populations may have contributed to our lack of statistical replication, despite the functional relevance of the *NFKB1* variants demonstrated by our functional assays. Our study expands the understanding of pharmacogenetic analyses in racially/ethnically diverse populations and advances the foundation for precision medicine in at-risk and understudied minority populations.

**AUTHOR SUMMARY:** Asthma is the most common chronic disease among children. Albuterol, a bronchodilator medication, is the first-line therapy for asthma treatment worldwide. In the U.S., asthma prevalence is the highest among Puerto Ricans, intermediate among African Americans and lowest in Whites and Mexicans. Asthma disparities extend to mortality, which is four- to five-fold higher in Puerto Ricans and African Americans compared to Mexicans [1]. Puerto Ricans and African Americans, the populations with the highest asthma prevalence and death rate, also have the lowest albuterol bronchodilator drug response (BDR). We conducted the largest pharmacogenetic study using whole genome sequencing data from 1,441 minority children with asthma who had extremely high or low albuterol bronchodilator drug response. We identified population-specific and shared pharmacogenetic variants associated with BDR. Our findings help inform the direction of future development of asthma medications and our study advances the foundation of precision medicine for at-risk, yet understudied, racially/ethnically diverse populations.

## INTRODUCTION

Asthma is a chronic inflammatory disorder of the airways characterized by recurrent respiratory symptoms and reversible airway obstruction. Asthma affects 5% of the world population [2] and is the most common chronic disease among children [3, 4]. In the United States (U.S.), asthma is the most racially disparate health condition among common diseases [5, 6]. Specifically, U.S. asthma prevalence is highest among Puerto Ricans (36.5%), intermediate among African Americans (13.0%) and European Americans (12.1%), and lowest among Mexican Americans (7.5%) [7]. These disparities also extend to asthma mortality, which is four- to fivefold higher in Puerto Ricans and African Americans compared to Whites and Mexican Americans [1].

Current asthma guidelines recommend inhaled β2-agonists (e.g., albuterol) for treatment of acute asthma symptoms. Albuterol is a short-acting β2-adrenergic receptor (β2AR) agonist, and it produces bronchodilation by causing rapid smooth muscle relaxation in the airways. Albuterol is the most commonly prescribed asthma medication in the world and is the mainstay of acute asthma management across all ethnic groups [8, 9]. Among low income and minority populations in the U.S., albuterol is often the only medication used for asthma regardless of asthma severity [10, 11]. Response to albuterol is quantified based on bronchodilator drug response (BDR) using spirometry. We and others have demonstrated that there is significant variability in BDR among individuals and between populations [12, 13]. Specifically, the populations with the highest asthma prevalence and mortality also have the lowest drug response to albuterol: Puerto Rican and African American children have significantly lower BDR than Whites and Mexican American children [13, 14]. This variation in drug response across racial/ethnic groups may contribute to the observed disparities in asthma morbidity and mortality [15–19].

BDR is a complex trait, influenced by environmental and genetic factors, with heritability estimates ranging from 47% to 92% [20–22]. Genome-wide association studies (GWAS) have identified several common single nucleotide polymorphisms (SNPs) associated with BDR in populations of European descent [23–25]. To date, only one GWAS of BDR has been conducted among African Americans [26]. While that study identified a novel BDR-associated locus, it did not replicate known associations discovered in populations of European descent, suggesting that BDR may be determined in part by population-specific variants. Our previous study of genetic predictors of BDR in Latino populations identified a significant contribution of population-specific rare variants to BDR [27].

GWAS were designed to identify common variants associated with disease through the use of genotyping arrays that relied on linkage disequilibrium to tag/represent variants not explicitly genotyped on the array itself. Early GWAS arrays were optimized for performance in populations of European origin and lacked the ability to capture race-/ethnic-specific genetic variation due to differences in linkage disequilibrium (LD) across racially/ethnically diverse populations [28]. Recent generations of arrays have attempted to tailor genotyping panels for major HapMap populations (Affymetrix Axiom^®^ World Arrays [29]), or to include population-specific and trans-ethnic tag SNPs to statistically infer genotypes not directly captured in diverse populations (Illumina Infinium^®^ Multi-Ethnic Genotyping Array [30]). However, imputation accuracy decreases significantly with variant frequency [31, 32], making it difficult to use genotyping arrays to study rareand/or population-specific variants.

The most striking weakness of GWAS is the inability to adequately capture rare variation. Whole exome sequencing (WES) and other forms of targeted sequencing were developed to address the inability of genotyping arrays to capture rare variation. WES only allows for the capture of common and rare variants within coding and flanking regions. Studies have shown that a large number of variants associated with complex disease lie within non-coding regions of the genome (reviewed in Zhang and Lupski, 2015 [33]). Additionally, the target capture procedures result in uneven sequence coverage, limiting the reliability of SNP calling for loci close to the boundary of targeted regions. WES also has limited usage for the detection of structural variation, which depends heavily on uniform coverage across the genome.

Whole genome sequencing (WGS) is the ideal technology for identifying disease-causing variants that are rare and/or population-specific. Unlike GWAS genotyping arrays or targeted sequencing technologies, WGS allows the detection of common and rare variants in coding and non-coding regions. WGS is the only technology capable of a truly comprehensive and agnostic evaluation of genetic sequence variation in the context of complex disease. The persistent lack of large-scale genetic studies conducted in populations of non-European descent further exacerbates racial/ethnic disparities in clinical and biomedical research [34–36]. The application of WGS to the evaluation of genetic factors within a racially/ethnically diverse study population is a necessary step toward eliminating health disparities in BDR and other complex phenotypes.

In this study, we performed WGS on 1,441 minority children with asthma from the tails of the BDR distribution (S1 Fig). Our study included high and low drug responders from three ethnic groups: Puerto Ricans (PR) (n=483), Mexicans (MX) (n=483), and African Americans (AF) (n=475). An overview of the subject selection process and main analyses performed in this study is presented in **Fig 1**. We identified multiple BDR-associated common and rare variants that are population-specific or shared among populations. This study is part of the National Heart, Lung, and Blood Institute’s Trans-Omics for Precision Medicine Whole Genome Sequencing (TOPMed) program and represents the largest WGS study thus far to investigate genetic variants important for bronchodilator drug response in racially and ethnically diverse children with asthma.

**Fig 1.**
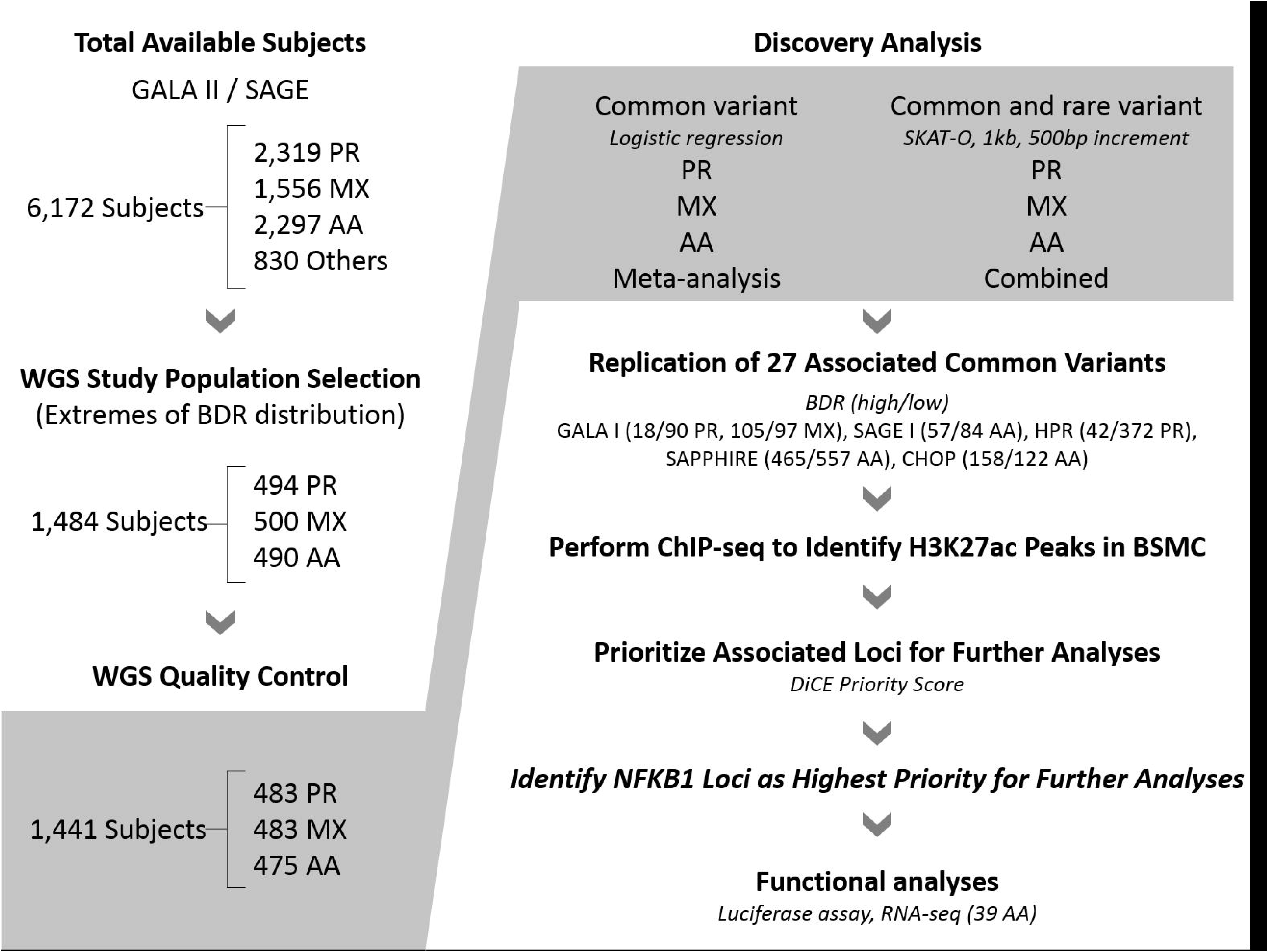
An overview of the main analyses performed in the current study. More detailed descriptions of the discovery and replication cohort demographics and analyses performed for common and rare variant analysis can be found in Methods.

## RESULTS

### Descriptive characteristics of study subjects

Descriptive characteristics for all study subjects (N=1,441, including 483 Puerto Ricans, 483 Mexicans and 475 African Americans) are summarized in Table 1. Covariates and demographic variables were assessed for significant differences between high and low drug responders for each racial/ethnic group. Significant differences were found for age (Mexicans, p<0.001), baseline lung function (pre-FEV_1_ % predicted, p<0.001), total Immunoglobulin E (tIgE, p < 0.001), and atopy. Baseline lung function (pre-FEV_1_ % predicted) was defined as the percentage of observed FEV_1_ relative to the expected population average FEV_1_ estimated using the Hankinson lung function prediction equations [37].

**Table 1.** Study Population Description (N=1,441).

| Descriptive Statistics |  | Puerto Ricans (N=483) |  |  | Mexicans (N=483) |  |  | African Americans (N=475) |  |  |
| --- | --- | --- | --- | --- | --- | --- | --- | --- | --- | --- |
|  |  | High BDR | Low BDR | P | High BDR | Low BDR | P | High BDR | Low BDR | P |
| Number of Subjects |  | 239 | 244 | - | 243 | 240 | - | 233 | 242 | - |
| Percent Male |  | 53.6% | 53.3% | 1.0 | 60.1% | 52.1% | 0.08 | 55.4% | 47.9% | 0.12 |
| Median Age, yr (IQR) |  | 11.6<br>(9.7 - 14.8) | 12.2<br>(10.1 - 15.2) | 0.18 | 11.7<br>(9.6 - 14.0) | 13.3<br>(10.6 - 16.0) | <0.001 | 13.8<br>(11.0 - 16.8) | 13.8<br>(10.9 - 17.1) | 0.48 |
| Mean Global Ancestry Proportions | AFR | 0.24 | 0.22 | 0.44 | 0.05 | 0.05 | 0.37 | 0.79 | 0.79 | 0.80 |
|  | EUR | 0.63 | 0.64 | 0.27 | 0.37 | 0.36 | 0.84 | 0.19 | 0.20 | 0.70 |
|  | NAM | 0.13 | 0.13 | 0.93 | 0.58 | 0.59 | 0.90 | 0.02 | 0.02 | 0.90 |
| BMI Category, N | Obese | 76 | 67 | 0.32 | 100 | 96 | 0.85 | 82 | 83 | 0.85 |
|  | Non-Obese | 163 | 177 |  | 143 | 144 |  | 151 | 159 |  |
| Pre-FEV<sub>1</sub> % Predicted, N | < 80% | 149 | 56 | <0.001 | 43 | 7 | <0.001 | 47 | 6 | <0.001 |
|  | ≥ 80% | 90 | 188 |  | 200 | 233 |  | 186 | 236 |  |
| Median ΔFEV<sub>1</sub>, % (IQR) |  | 21.2<br>(18.2 - 25.7) | 5.0<br>(2.9 - 6.3) | - | 12.7<br>(10.3 - 16.8) | 3.6<br>(2.0 - 4.9) | - | 15.5<br>(13.3 - 20.3) | 3.3<br>(2.0 - 4.4) | - |
| Median tIgE, mL (IQR) |  | 407.5<br>(126.8-952.8) | 191.9<br>(50.5-542.2) | <0.001 | 247.5<br>(64.2-817.0) | 105.7<br>(35.4-332.0) | <0.001 | 281.6<br>(97.3-552.4) | 128.8<br>(36.6-351.3) | <0.001 |
| Atopy, N |  | 177 | 118 | <0.001 | 155 | 117 | <0.001 | 139 | 111 | <0.001 |

**BDR**: bronchodilator drug response. IQR: interquartile range. **Pre-FEV_1_ % predicted**: percentage of measured FEV_1_ relative to predicted FEV_1_ estimated by the Hankinson lung function prediction equations prior to administration to albuterol. **ΔFEV_1_**: a quantitative measure of BDR, measured as the percent change in baseline FEV_1_ after administration of albuterol. High and low drug responders were chosen from the extremes of the BDR (ΔFEV_1_) distribution. **tIgE**: measure of total Immunoglobulin E from serum in milliliters. **Atopy**: tIgE measurement greater than or equal to 100. Table 1. Study Population Description (N=1,441).

We estimated genetic ancestry for all participants (see **Methods**) and found that the major ancestry proportions in Puerto Ricans, Mexicans and African Americans are European, Native American and African ancestries, respectively (**Table 1, S2 Fig**). Analysis of genetic substructure of the three admixed populations by principal component analysis (PCA) demonstrated that the three populations displayed the characteristic spectrum of ancestry found in admixed populations (**S3 Fig**).

### Variant summary statistics

Genetic variant summary statistics revealed that the average number of variants by population corresponded to the proportion of African ancestry: the most variants were found among African Americans, followed by Puerto Ricans and Mexicans (**Fig 2a, Table 2**).

**Fig 2.**
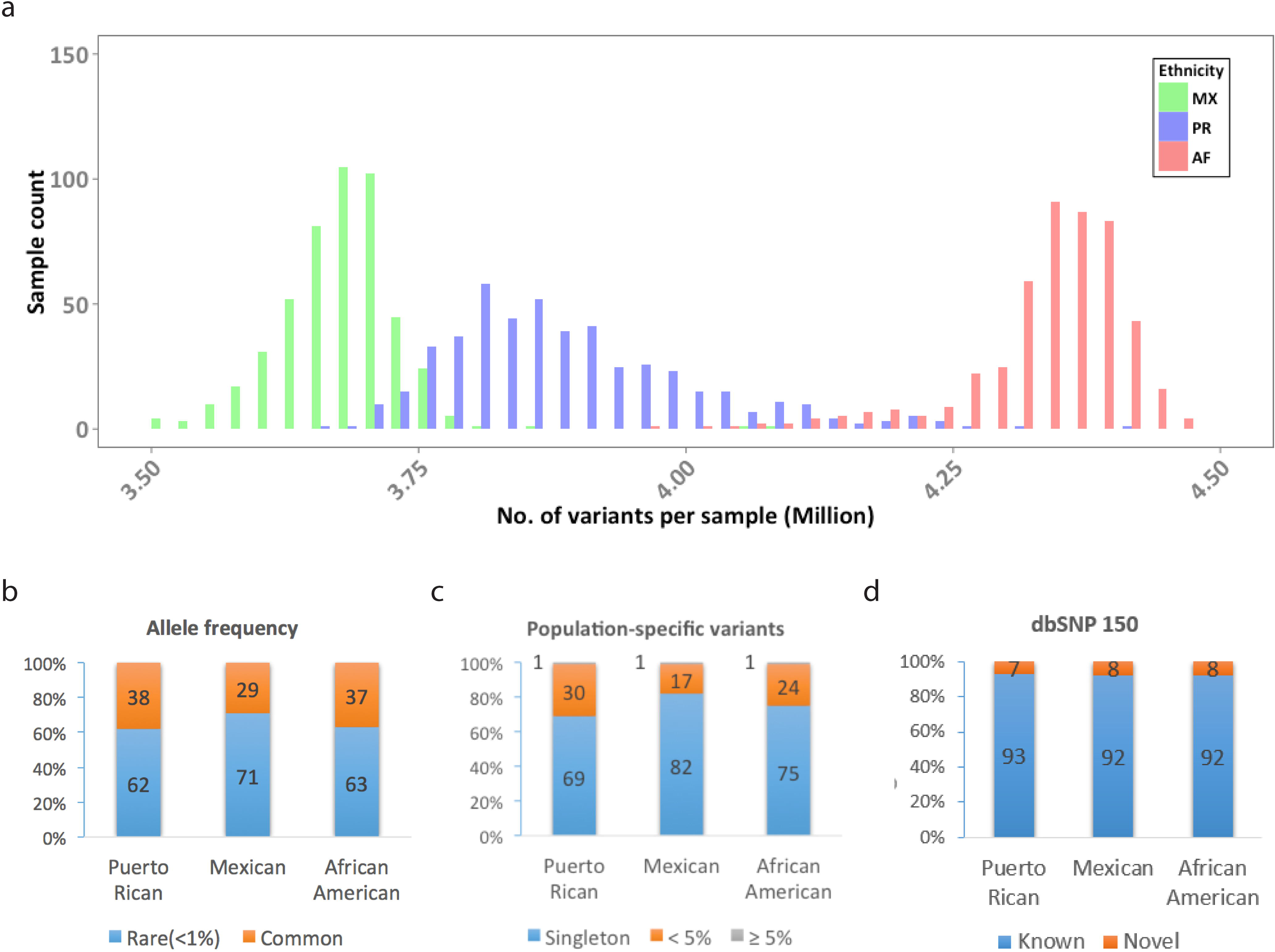
**(a)** Number of variants per sample. The bin size is 0.025M variants. **(b)** Allele frequency of biallelic SNPs (relative to GRCh37). **(c)** Allele frequency of population-specific biallelic SNPs. **(d)** Novel biallelic SNPs based on dbSNP build150.

**Table 2.** Summary statistics of variants

|  | Puerto Ricans | Mexicans | African Americans |
| --- | --- | --- | --- |
| **Range in variants, per individual (mean)** |  |  |  |
| **All** | 3.7M – 4.4 M<br>(4.0 M) | 3.5M – 4.1M<br>(3.7M) | 4.0M – 4.5M<br>(4.3M) |
| SNPs | 3.4M – 4.0 M<br>(3.6M) | 3.2M - 3.7M<br>(3.4M) | 3.6M – 4.1M<br>(4.0M) |
| Indels | 284,067 – 344,493<br>(305,618) | 272,635 – 321,778<br>(289,096) | 311,997 – 354,646<br>(339,570) |
| Others* | 21,233 – 29,177<br>(24,728) | 19,703 – 26,772<br>(23,190) | 22,014 – 30,993<br>(27,911) |
| **No. of biallelic SNPs, union\*\*** | 29.2M | 28.1M | 36.3M |
| **By allele frequency of data** |  |  |  |
| Rare (< 1%) | 18,169,292 (62%) | 20,029,291 (71%) | 22,847,938 (63%) |
| Common (≥ 1%) | 11,007,125 (38%) | 8,092,157 (29%) | 13,459,004 (37%) |
| **Population-specific\*\*\*** | 6,680,909 | 9,687,651 | 14,114,142 |
| Singleton | 4,616,383 (69%) | 7,983,950 (82%) | 10,574,879 (75%) |
| < 5% | 6,676,286 (> 99%) | 9,675,271 (> 99%) | 14,096,844 (> 99%) |
| ≥ 5% | 4,623 (< 1%) | 12,380 (< 1%) | 17,298 (< 1%) |
| **By novelty (dbSNP150)** |  |  |  |
| Known | 27,034,179 (93%) | 25,911,801 (92%) | 33,540,387 (92%) |
| Novel | 2,142,238 (7%) | 2,209,647 (8%) | 2,766,555 (8%) |
| **By protein impact\*\*\*\*** |  |  |  |
| Coding | 284,269 (1%) | 289,055 (1%) | 362,823 (1%) |
| *Nonsynonymous* | 157,541 | 164,312 | 203,029 |
| *Stopgain / stoploss* | 3,194 | 3,531 | 4,158 |
| *Splicing* | 2,111 | 2,214 | 2,620 |
| *Other coding* | 121,423 | 118,998 | 153,016 |
| Noncoding | 28,833,461 (99%) | 27,769,434 (99%) | 35,878,723 (99%) |
| Not annotated^ | 58,687 (< 1%) | 62,959 (< 1%) | 65,396 (< 1%) |

* Includes multi-nucleotide polymorphism (MNPs), complex, symbolic and mixed variants as defined by GATK VariantEval.

** Biallelic SNPs with less than 10% genotype missingness per population were included.

*** Biallelic SNPs that are present in only one of the three studied populations.

**** Other coding variants include those annotated as exonic and synonymous in ANNOVAR.

^ Not annotated: biallelic SNPs not included in the annotation pipeline because they are not present in TOPMed freeze 2 and 3 data releases.

The majority of observed variants (>90%) were SNPs. The union of biallelic SNPs from all individuals in each population varied from 28.1M among Mexicans, 29.2M among Puerto Ricans to 36.3M among African Americans. Approximately 65% of biallelic SNPs were rare (non-reference allele frequency < 1%, **Fig 2b, Table 2**). Biallelic SNPs that were population-specific (i.e., SNPs found in only one population) accounted for 23% (6.68M / 29.2M in Puerto Ricans) to 39% (14.1M / 36.3M in African Americans) of the biallelic SNPs observed in each population. Over 99% of the population-specific SNPs had a non-reference allele frequency less than 5% and the majority of these population-specific SNPs (69% to 82%) were also singletons (**Fig 2c, Table 2**). Based on dbSNP build 150, an average of 8% of biallelic SNPs were novel (**Fig 2d, Table 2**).

In all three populations, 99% of the biallelic SNPs were observed in noncoding regions. Based on the Combined Annotation Dependent Depletion (CADD) score [38], which estimates the deleteriousness of a variant, over 99% of highly deleterious biallelic SNPs (CADD score ≥ 25) were observed in coding regions, regardless of ethnicity (**Table 3**). This may be due to the relatively limited availability of functional annotations as positive training data for CADD to estimate deleteriousness in non-coding regions [39]. The percentage of singletons in these highly deleterious biallelic SNPs varied from 51% (Puerto Ricans) to 70% (Mexicans).

**Table 3.** CADD score summary statistics of biallelic SNPs*.

| Population | CADD score | No. of SNPs* | % Coding | % Singletons |
| --- | --- | --- | --- | --- |
| Puerto Ricans | 0-9 | 26,936,285 | 0.5% | 29% |
|  | 10-19 | 2,027,392 | 4% | 32% |
|  | 20-24 | 123,481 | 32% | 37% |
|  | ≥ 25 | 30,572 | >99% | 51% |
| Mexicans | 0-9 | 25,932,006 | 0.5% | 44% |
|  | 10-19 | 1,967,096 | 4% | 49% |
|  | 20-24 | 124,173 | 34% | 55% |
|  | ≥ 25 | 35,214 | >99% | 70% |
| African Americans | 0-9 | 33,468,245 | 0.5% | 36% |
|  | 10-19 | 2,573,523 | 4% | 40% |
|  | 20-24 | 158,967 | 32% | 46% |
|  | ≥ 25 | 40,811 | >99% | 64% |

*Biallelic SNPs with less than 10% genotype missingness per population were included.

### BDR association testing with common variants

We performed genome-wide association testing of common variants with BDR (dichotomized as high/low drug responders from the extremes of the BDR distribution) for each population, adjusting by age, sex, body mass index (BMI) categories, and the first ten principal components (PCs) (see **Methods** section “Single locus BDR association testing on common variants” for rationale on including these covariates). We then performed a trans-ethnic meta-analysis on these results across all three populations. For all three populations, 94% of all common variants tested are known variants annotated in dbSNP Build 150 (S1 Table).

A universal *p*-value threshold of 5 × 10^-8^ is often used to determine significance in GWAS. This statistical threshold was calculated based on Bonferroni correction under the assumption of 1,000,000 independent tests using patterns of linkage disequilibrium based primarily on individuals of European descent and has been shown to be non-generalizable for WGS studies or genetic studies of non-European populations in general [40–42]. The number of independent tests varies by LD patterns, which in turn vary by race/ethnicity [41]. We calculated the effective number of independent tests for each population and for our trans-ethnic metaanalysis, and generated racially/ethnically adjusted genome-wide significance thresholds for each population (see **Methods**). Population-specific genome-wide significance thresholds after correcting for the number of effective tests (adjusted genome-wide significance level) were 1.57 × 10^-7^ for Puerto Ricans, 2.42 × 10^-7^ for Mexicans, and 9.59 × 10^-8^ for African Americans (see **Methods**). These numbers are highly concordant with WGS significance thresholds derived from the African American (ASW), Mexican (MXL), and Puerto Rican (PUR) 1000 Genomes sequencing data [41]. The adjusted genome-wide significance level for our transethnic meta-analysis was 3.53 × 10^-7^. Significance thresholds for discovery analyses in genome-wide association studies can often produce false negative results [43–46]. To minimize Type II error, suggestive associations are often included in replication and functional validation studies. We identified suggestive associations based on the following widely used formula: 1/(effective number of tests) [43–46].

While no significant associations were identified from the population-specific analyses (**S8 Fig**), our trans-ethnic meta-analysis identified ten unique loci (represented by 27 SNPs) significantly (*p* < 3.53 x 10^-7^) or suggestively (*p* < 7.06 × 10^-6^) associated with BDR status (**Fig 3a, Table 4, S2 Table**). We annotated all 27 SNPs by performing a thorough bioinformatics search in ENCODE, NHGRI-EBL GWAS Catalog and PubMed databases. Their previously reported lung-related phenotype associations and functional annotations are reported in **S4 Table** and **S9 Table**. Two SNPs, rs17834628 and rs35661809, located on chromosome 5 were significantly associated with BDR (*p* = 1.18 x 10^-8^ and 3.33 x 10^-8^); additionally, population-specific analyses show that the direction of effect for these two variants is concordant across all three populations (**Fig 3b, S3 Table**). **Fig 3c** displays a LocusZoom plot of rs17834628 with 400 kb flanking regions. Three of the 27 identified SNPs were located within genes. Specifically, two SNPs are located in the third and fifth introns of *NFKB1* (rs28450894 and rs4648006), and a third SNP, rs16995064, mapped to intron 7 of *PLCB1* (**Table 4**). Among the *NFKB1* SNPs, the low BDR-associated T allele of rs28450894 is found predominantly among African populations (minor allele frequency [MAF] 8.8% – 28.7%), followed by European populations (MAF 3.7% – 7.6%) and Puerto Ricans (MAF 6.2%), and is relatively rare in Mexicans (MAF 1.5%) based on 1000 Genomes data (**S6 Fig**). Combined, these 27 SNPs explain 23%, 16%, and 18% of the variation in BDR status in Puerto Ricans, Mexicans, and African Americans, respectively (**S5 Table**).

**Fig 3.**
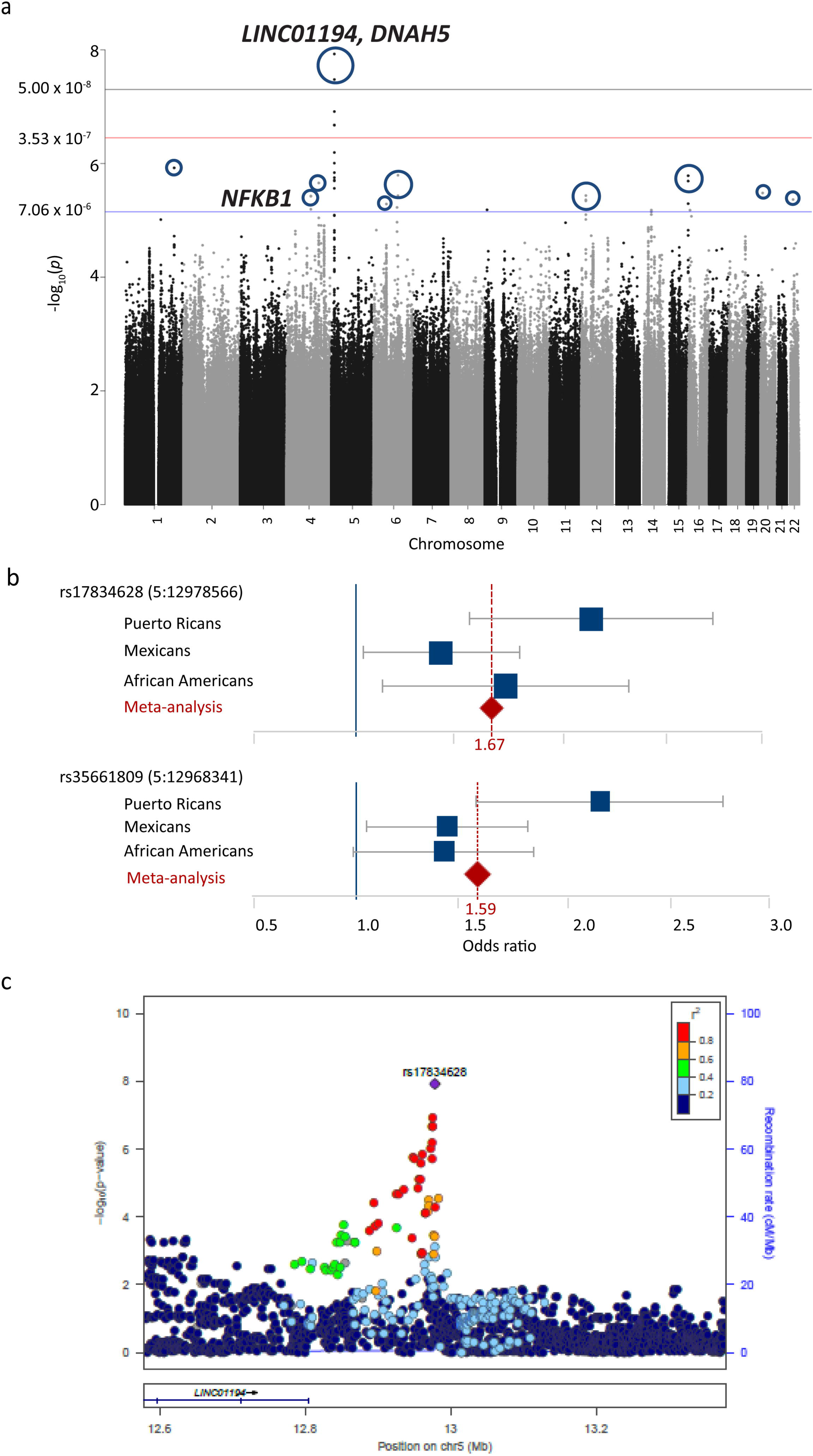
**(a)** Manhattan plot of the trans-ethnic meta-analysis of single locus BDR association testing. Top ten BDR-associated loci are circled. Black line indicates universal genome-wide significance threshold (5.00 x 10^-8^), red line indicates adjusted genome-wide significance threshold (3.53 x 10^-7^), and blue line indicates suggestive significance threshold (7.06 x 10^-6^). **(b)** Forest plot of the population-specific and joint effect of the two most significantly associated SNPs, rs17834628 and rs35661809. The R^2^ between these two SNPs is 0.93 in Puerto Ricans, 0.96 in Mexicans and 0.66 in African Americans. **(c)** The most significantly associated SNP (rs17834628) is plotted together with 400kb flanking regions on either side. Color of the dots shows the LD of each SNP with rs17834628 based on the 1000 Genomes Nov 2014 AMR population. Multiple SNPs in high LD (R^2^ > 0.8, red) reached a suggestive significance level.

**Table 4.**
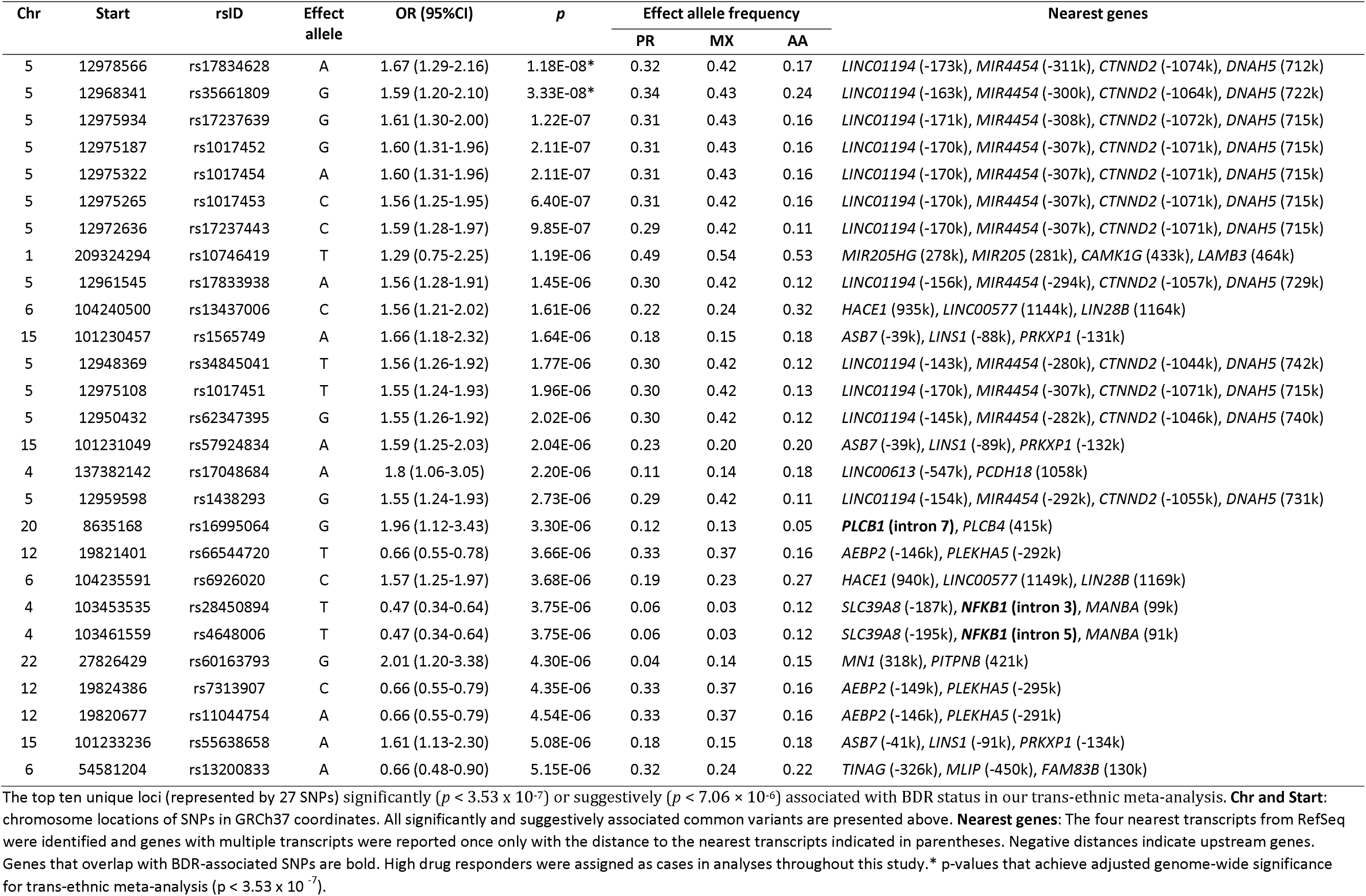
Results from trans-ethnic BDR association tests for common variants.

The top ten unique loci (represented by 27 SNPs) significantly (p < 3.53 x 10^-7^) or suggestively (p < 7.06 × 10^-6^) associated with BDR status in our trans-ethnic meta-analysis. **Chr and Start**: chromosome locations of SNPs in GRCh37 coordinates. All significantly and suggestively associated common variants are presented above. **Nearest genes**: The four nearest transcripts from RefSeq were identified and genes with multiple transcripts were reported once only with the distance to the nearest transcripts indicated in parentheses. Negative distances indicate upstream genes. Genes that overlap with BDR-associated SNPs are bold. High drug responders were assigned as cases in analyses throughout this study.* *p*-values that achieve adjusted genome-wide significance for trans-ethnic meta-analysis (p < 3.53 x 10).

It has been shown that functionally relevant variants do not always display the lowest *p*-values in association studies [47]. To avoid false negative results, it is strongly suggested that replication analyses for two-stage genome-wide studies [43–46]. Therefore, we included all 27 SNPs in replication analyses we performed separately and via meta-analysis in five independent populations (GALA I, SAGE I, HPR, SAPPHIRE and CHOP) (S6-S7 Table). None of the 27 SNPs were significantly associated with BDR status in our replication analyses (S6-S7 Table). It is important to note that our largest replication cohort (SAPPHIRE) did not include children (median age = 30 and 28 for high and low BDR groups, respectively, S6 Table). All other replication populations included less than 500 individuals per study (S6 Table). It is well known that there are age-specific associations with asthma and asthma-related phenotypes [48, 49]. It is unclear whether the same is true for BDR. It is possible that the overrepresentation of adult patients in our available replication populations may explain, in part, our lack of replication.

In addition to performing WGS association analyses to identify genetic variants associated with variation in BDR, we also performed H3K27ac chromatin immunoprecipitation sequencing analysis (ChIP-seq) experiments in primary bronchial smooth muscle cells (BSMCs) to identify potential regulatory regions marked by H3K27ac peaks. Albuterol’s mechanism of action involves binding with the β2-adrenergic receptor in bronchial smooth muscle cells causing rapid onset of airway tissue relaxation and bronchodilation. BSMCs are therefore considered one of the most relevant cell types for molecular studies of BDR [50]. We observed two H3K27ac ChIP-seq signals that overlapped with variants in moderate to high LD (R^2^ = 0.47 to 0.82) with two *NFKB1* SNPs (rs28450894 and rs4648006) we identified through trans-ethnic meta-analysis, implying that these variants may have regulatory functions in BSMC (**S5 Fig a, S8 Table**).

The gold standard for identifying true signals in genetic association studies is to use *p*-values from a primary and/or replication study to prioritize variants for further investigation. The use of *p*-values as the sole metric for prioritization is problematic for three reasons: (1) the p-value statistic is dependent on sample size and effect magnitude, (2) *p*-values do not incorporate biological knowledge, and (3) one cannot use *p*-values to distinguish between true association signals and noise of the same magnitude [47, 51, 52]. Instead of relying solely on *p*-values, we applied the Diverse Convergent Evidence (DiCE) [53] approach to prioritize each of the 27

BDR-associated SNPs from our trans-ethnic meta-analysis for inclusion in further function analyses (**S15 Table, S4 Fig**). After integrating information from our WGS analysis, publicly available bioinformatics data, and ChIP-Seq experiments in BSMCs, the *NFKB1* locus had the highest DiCE evidence score, indicating that this locus had the strongest evidence of functional relevance to BDR variation (**S4 Fig**). Therefore, all further functional experiments were focused on variants within this locus.

### Functional assays on the *NFKB1* Locus

Both H3K27ac ChIP-seq regions that overlapped with the BDR-associated *NFKB1* locus were tested for enhancer activity using luciferase enhancer assays. The sequences of these two *NFKB1* intronic regions were cloned into a pGL4.23 enhancer assay vector (Promega, Madison, WI, USA), which contains a minimal promoter and a luciferase reporter gene. The pGL4.23 vector with the viral SV40 promoter was used as a positive control, and the pGL4.23 empty vector as a negative control. All constructs were tested for their enhancer activity in BSMCs.

One enhancer, *NFKB1* Region 2, showed significantly increased enhancer activity over empty vector (2.24-fold increase, p = 8.70 x 10^-6^, unpaired t-test; **S5 Fig b**). Given the relevance of *NFKB1* in immune pathways and asthma, we also performed RNA sequencing (RNA-seq) experiments to verify whether the identified intronic *NFKB1* SNPs regulate gene expression of neighboring genes. Among genes within 1Mb of rs28450894 meeting expression reliability cutoffs (see Methods), we found that the low BDR-associated T allele of rs28450894 is significantly associated with decreased expression of SLC39A8 in blood (**S7 Fig**, p = 0.0066, FDR-adjusted p = 0.0856, log2(β) = -0.327).

We observed that two known BDR candidate genes, ADCY9 and CRHR2, which achieved replication in a previous GWAS of BDR performed in the full GALA II population but did not replicate in the current study (S10 Table) [27]. In the previous study, GWAS array data, supplemented by imputation, were used to evaluate genetic associations with BDR measured as a continuous trait. To determine whether the discrepancy between findings was due to data type (imputed array-based vs. WGS-based) or study design (continuous trait vs. extreme phenotype), the common variant analysis in the current analysis was repeated among the subset of samples that had array-based and WGS data (n = 1,414 out of 1,441). Based on the top 1,000 BDR-associated SNPs from the current common variant analysis, there was high correlation between association *p*-values generated from imputed array-based and WGS-based genotypes (Spearman correlation = 1.0), suggesting that data type is not the cause of the observed discrepancy (**S9 Fig a**).

Nearly all SNPs with high imputation R^2^ exhibited high genotype concordance between array-based and WGS-based genotypes, confirming high imputation quality for most common SNPs (≥ 99.7%). (**S9 Fig b and c**). We also performed linear regression to analyze BDR (ΔFEV_1_) as a continuous trait using imputed array-based data. The most significantly associated SNP identified in the trans-ethnic metaanalysis using extreme phenotype analysis displayed the same direction of effect as analyzing BDR as a continuous trait (OR=1.67 in extreme phenotype analysis and β=0.51 in continuous analysis). These observations indicate that the discrepancy between findings may be due to differences in statistical power afforded by the different study designs (continuous trait vs. extreme phenotype). For common variant analyses, dichotomization of a continuous outcome results in a loss of statistical power [54–57]. For example, a population of 2,000 individuals has 80% power to identify moderate genetic associations (β=0.3) for common variants with minor allele frequencies ≥ 0.05 when the outcome is continuous. If this population were re-analyzed after dichotomizing the continuous outcome at the median (cases=1,000, controls=1,000), power would be reduced to 62%. The opposite effect is observed in rare variant analyses. The extreme phenotype study design is a specific type of dichotomous outcome study design that has been shown to increase power and the probability of identifying functional rare variants [56–58]. It should also be noted that the previously published results were discovered in one population (Puerto Ricans), whereas the results from our trans-ethnic meta-analysis describe associations that are conserved across three populations (Puerto Ricans, Mexicans, and African Americans).

### BDR association testing using common and rare variants

We tested the combined effects of common and rare variants on BDR using SKAT-O [59] to examine variants in 1kb sliding windows, which moved across the genome in 500bp increments. The same covariates used for common variant association testing were applied.

After identifying the effective number of tests and adjusting for multiple comparisons on each population separately (see Methods), we identified three population-specific loci associated with BDR at genome-wide significance levels; two were found in Mexicans on chromosome 1 and chromosome 11, and one in African Americans on chromosome 19 (**Fig 4a-c, Table 5, S11 Table**).

**Fig 4.**
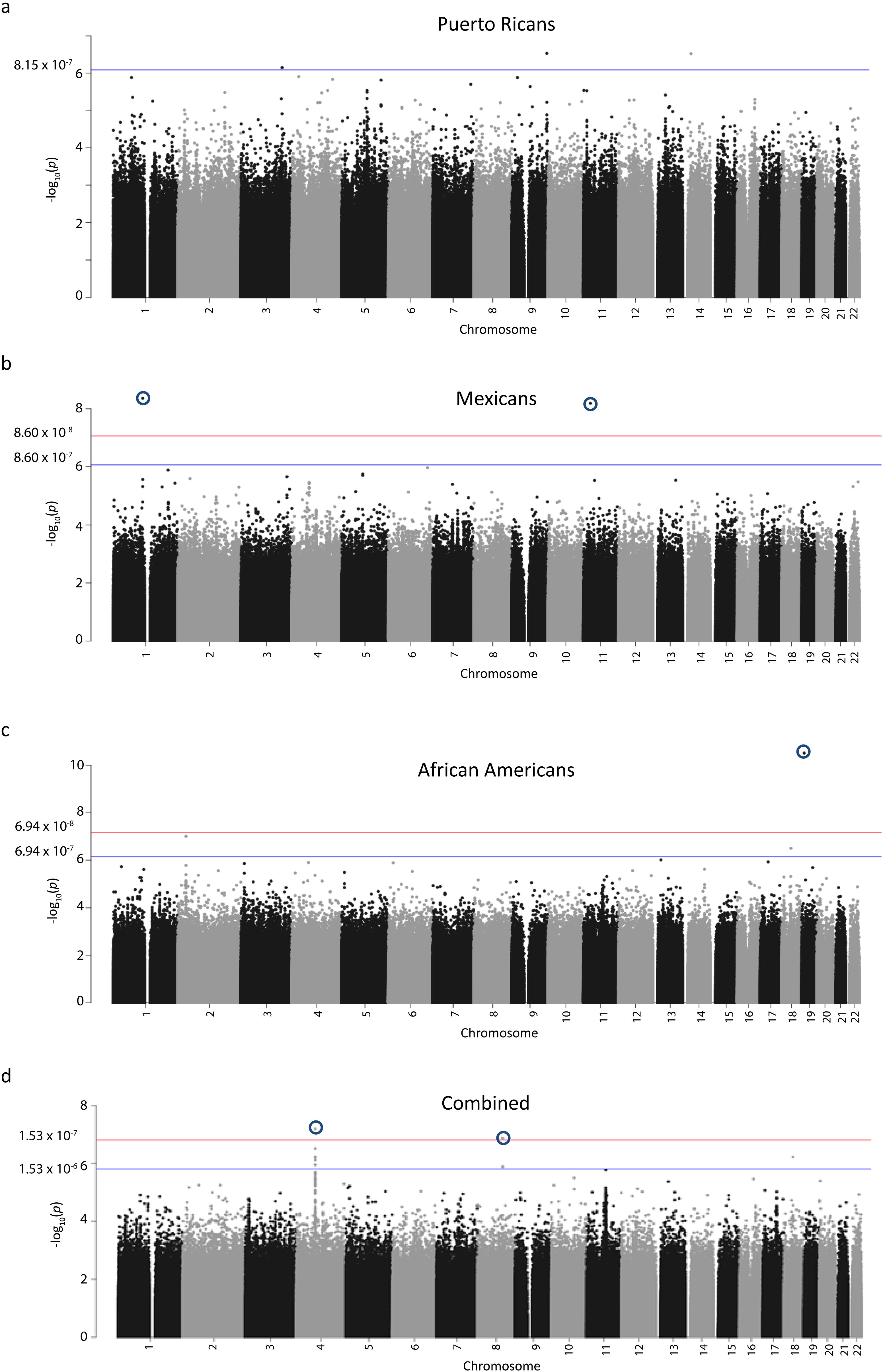
Manhattan plot of SKAT-O analysis of biallelic common and rare SNPs grouped by 1kb windows sliding across chromosome 1 to 22 in **(a)** Puerto Ricans **(b)** African Americans, **(c)** Mexicans, and **(d)** all populations combined. Bonferroni-corrected genome-wide and suggestive significance levels are marked by red and blue lines, respectively.

**Table 5.** Results from association testing on combined effects of common and rare variants on BDR Chr, start and stop: GRCh37 chromosome coordinates; AA: African Americans; MX: Mexicans; Combined: all individuals in all three populations. nCommon and nRare: number of common and rare variants, respectively. Nearest genes: The four nearest transcripts from RefSeq were identified and genes with multiple transcripts were reported once only with the distance to the nearest transcripts indicated in parentheses. Negative distances indicate upstream genes. Genes that overlap with BDR-associated SNPs are bold.

| Chr | Start | Stop | *p* | Population | nCommon | nRare | Nearest genes |
| --- | --- | --- | --- | --- | --- | --- | --- |
| 1 | 114177000 | 114178000 | 4.40E-09 | MX | 2 | 1 | **MAGI3 (intron 9)**, *PHTF1* (62k), *RSBN1* (126k) |
| 11 | 27507000 | 27508000 | 6.59E-09 | MX | 2 | 3 | *LOC105376671* (-3k), *LGR4* (-13k), *LIN7C* (8k) |
| 19 | 10424000 | 10425000 | 3.12E-11 | AA | 1 | 2 | *ZGLP1* (-4k), *ICAM5* (-17k), **FDX1L (intron 3)**, *RAVER1* (2k) |
| 4 | 73478000 | 73479000 | 6.25E-08 | Combined | 10 | 23 | *ADAMTS3* (-43k), *COX18* (441k) |
| 8 | 97926000 | 97927000 | 1.32E-08 | Combined | 3 | 13 | *SDC2* (-302k), **CPQ (intron 4)**, *LOC101927066* (37k), *TSPYL5* (359k) |

We also performed association testing across all three populations in a single analysis. Pooling subjects increased the sample size and thereby maximized the power of the SKAT-O association test. To minimize any potential effect of confounding by population substructure, association testing also included local genetic ancestry, defined as the proportions of Native American and African ancestries for the window under testing. Two loci on chromosomes 4 and 8 were found to be genome-wide significant (p < 1.53 × 10^-7^) (**Fig 4d, Table 5**). A total of 60 variants were identified from all SKAT-O regions reported in Table 5. Six of the 60 variants were located within predicted regulatory regions (S12 Table). Specifically, three variants located on chromosome 11 identified in Mexicans overlap with a CTCF (transcriptional repressor) binding site and comprise a chromatin insulator region. The five regions identified in our combined and population-specific SKAT-O analyses independently explained 4% to 8% of the variation in BDR in their respective populations (**Table 5, S5 Table**).

We examined alternative grouping strategies for rare variants, including grouping by (1) genes from transcription start to end sites with or without 50kb flanking regions, (2) transcription start site with 20kb flanking regions, and (3) H3K27ac ChIP-seq peaks in airway epithelial cells and airway smooth muscle cells. Association tests with these alternate grouping strategies identified no further significant associations.

## DISCUSSION

We identified population-specific and shared common and rare variants associated with bronchodilator drug response in three ethnically diverse populations of children with asthma. WGS, unlike GWAS genotyping arrays and targeted sequencing, provides comprehensive detection of common and rare variants in coding and non-coding regions. African Americans, Latinos, and other minorities have been dramatically underrepresented in GWAS [34–36]. Combined, the 27 variants identified from our common variant analyses (**Table 4**) explained 23%, 16%, and 18% of the variation in BDR in Puerto Ricans, Mexicans, and African Americans, respectively, after adjusting for clinical covariates (S5 Table). The five SKAT-O regions identified in our combined and population-specific analyses independently explained 4% to 8% of the BDR variation in their respective populations (**Table 5, S5 Table**). Our study represents an important investment from the NIH/NHLBI to include underrepresented populations in large whole genome sequencing efforts and to improve racial/ethnic diversity in clinical and biomedical research.

Our trans-ethnic common variants meta-analysis identified one locus on chromosome 5 that was associated with BDR at a genome-wide significance level (p < 5 x 10^-8^). The proximity of this BDR-associated locus to *DNAH5* and *LINC01194* is of particular interest. A SNP in *DNAH5* has been associated with total lung capacity in White subjects with chronic obstructive pulmonary disease [60]. In a separate GWAS, the *DNAH5*/*LINC01194* locus was reported among Europeans to be associated with levels of IgE [61, 62], a biomarker associated with asthma endotypes. Baseline lung function (FEV_1_) and total IgE levels are associated with asthma severity and can predispose an individual to lower bronchodilator drug responsiveness [13, 14, 63]. We found two *NFKB1* intronic variants on chromosome 4 associated with BDR at a suggestive significance level. The NFκB protein has a known role in allergic response, and various studies have demonstrated that the NFκB pathway is activated in patients with asthma, as reviewed by Edwards et al.[64].

ChIP-seq and functional enhancer assays in BSMCs suggest these *NFKB1* intronic variants may regulate expression of nearby genes. This was in fact supported by our RNA-seq data, which showed that individuals with the low BDR-associated T allele genotype displayed reduced expression of the neighboring gene SLC39A8, which has previously been found to be responsive to cytokine treatment in airway epithelial cells [65] and had reduced expression in mice with allergic airway inflammation [66]. Recent studies have also shown that SLC39A8 is unique among zinc transporters in that upregulation of SLC39A8 is sufficient to protect lung epithelium against TNF-α-induced cytotoxicity [67]. Additionally, the higher frequency of the low BDR-associated allele (T allele of rs28450894 in NFKB1) in African populations suggests that the low BDR-associated allele tracks with African ancestry. This may explain why admixed populations with higher proportions of African ancestry, i.e., African Americans and Puerto Ricans, have lower bronchodilator drug responsiveness [14], and by extension may shed light on the higher asthma morbidity and mortality in these populations.

Another intronic variant (chromosome 20, rs16995064, *PLCB1* intron 7) was associated with BDR at a suggestive significance level. *PLCB1* is highly relevant, as this gene has been reported to be differentially expressed in therapy-resistant childhood asthma compared to controlled persistent asthma or age-matched healthy control subjects in a Swedish cohort [68]. Functional studies also reported that silencing *PLCB1* inhibited the effect of lipopolysaccharide-induced endothelial cell inflammation through inhibiting expression of proinflammatory cytokines [69]. Further functional studies are necessary to establish the role of *NFKB1* and *PLCB1* on BDR.

Apart from assessing the individual effect of common variants on BDR, we also identified various combined effects of rare variants that were population-specific or shared across populations. While some of the nearest genes are uncharacterized or have no known functional relationship to BDR (*MAGI3, LOC105376671, LIN7C and CPQ*), there appears to be functional relevance for the locus between *ADAMTS3* and *COX18*. The *ADAMTS3* and *COX18* locus were associated with *β*-adrenergic responses in cardiovascular-related traits in mice [70]. This locus was significantly associated with cardiac atrial weight in mice treated with the *β* blocker atenolol; the association also replicated in mice treated with the *β* agonist isoproterenol. These findings suggest that SNPs found in this locus may modify *β* adrenergic signaling pathways in BDR. In the present study, we also identified BDR association with rare variants within the CPQ gene, which encodes a protein from the carboxypeptidase family. Although no previous BDR association has been identified for CPQ, another member of the carboxypeptidase family, carboxypeptidase A3 (CPA3), is known to be expressed at higher levels in the airway epithelium among subjects with TH2-high asthma [71, 72]. Further studies are necessary to determine the role of CPQ in BDR.

GWAS-based BDR-associated common variants in GALA II have previously been reported [27]. However, these variants did not replicate in the current study, likely due to different study designs between the previous and current investigations. The previous BDR GWAS used an array-based genotyping panel to examine children with asthma across the entire BDR spectrum, i.e., BDR was used as a continuous variable. In contrast, the current study sequenced the entire genome to investigate only the extremes of the BDR distribution (i.e., high and low drug responders). By repeating our current analysis using a subset of individuals who had array and WGS data, we confirmed that the major discrepancy between the two studies is due to study design instead of differences in data type. The contrast in results between GWAS and WGS due to differences in study design implies that varied study designs are necessary for a comprehensive understanding of variants associated with asthma-related phenotypes and drug response. Studying samples from the extreme tails of drug response distribution has been recognized as one of the success factors in the study design of pharmacogenomic GWAS [73]. Furthermore, it was recently demonstrated that the power gain from studying extreme phenotypes is much greater in analyses of rare variants compared to common variant studies [55]. Since cost is often a limiting factor for WGS studies, choosing an extreme phenotypic study design may be beneficial for the study of rare variants and the discovery of common variant associations that may otherwise be missed when sampling across the entire phenotypic spectrum.

We did not identify BDR-associated variants from β2AR signaling pathways. Instead, most of the BDR-associated genes identified in this study are related to lung function and allergic response, including total IgE levels and cytokine production in mast cells. This suggests that at least part of BDR may be due to the predisposition or intrinsic state of airway smooth muscle cells. Genetic variation may determine individuals’ intrinsic expression levels of candidate genes, which in turn determine whether their response to albuterol is beneficial.

A higher percentage of African ancestry often implies a higher degree of genetic variation [74]. Although Puerto Ricans have higher proportions of African ancestry than Mexicans (**Table 1**), they have fewer population-specific SNPs, an observation that is consistent with findings from our contributions to the 1000 Genomes Project [75] and our independent work. This is likely due to the fact that Puerto Ricans have gone through recent population bottlenecks [76]. We demonstrated that Puerto Ricans may be more genetically related than expected [76], suggesting that our current relatedness filters may be too conservative for Puerto Ricans.

Including admixed populations in whole genome sequencing studies has important scientific implications. First, it allows for discovery of genetic variation of multiple ancestral populations in a single study. Second, it is extremely useful to study admixed populations with ancestries that are currently underrepresented in existing genetic repositories. For example, the widely popular PCSK9 inhibitors used to treat hypercholesterolemia were discovered by studying the genetics of African Americans but the biology and final drug development have benefited all patients regardless of race/ethnicity [77]. Finally, studying admixed populations such as Mexicans will enhance the understanding of genetic variation in Native American ancestry, an area that is currently lacking in all major sequencing efforts.

Although an extensive effort was made to replicate the top BDR-associated variants, we were unable to replicate our results because few studies of non-European populations exist, as we and others have documented [34-36, 78]. Our efforts to perform replication of rare BDR-associated variants were further hindered by the lack of studies with whole genome sequencing data. These challenges highlight the need to include more racially/ethnically diverse populations in all clinical and biomedical research.

In an era of precision medicine, addressing questions about the impact of genetic factors on therapeutic drug response in globally diverse populations is essential for making precision medicine socially and scientifically precise [5]. This study advances our understanding of genetic analysis in admixed populations and may play an important role in advancing the foundation of precision medicine for understudied and racially and ethnically diverse populations.

## METHODS

### Data availability

TOPMed whole genome sequencing data are available to download by submitting a data access request through dbGaP. The dbGaP study accession numbers for GALA II and SAGE II are phs000920.v1.p1 and phs000921.v1.p1. WGS and array genotype data for each study are available through dbGaP under the same accession numbers.

### Study cohorts and sample details

This study examined a subset of subjects with asthma from the Study of African Americans, Asthma, Genes & Environments (SAGE II) [49, 79-81] and the Genes-Environments & Admixture in Latino Americans (GALA II) study [27]. SAGE II recruited African American subjects from the San Francisco Bay area. GALA II recruited Latino subjects from Puerto Rico and the mainland United States (Bronx, NY; Chicago, IL; Houston, TX; San Francisco Bay Area, CA). Ethnicity of the subjects was self-reported and all four of the participant’s biological grandparents must have reported the same ethnicity.

A total of 1,484 individuals from three ethnic groups (494 Puerto Ricans, 500 Mexicans and 490 African Americans), representing the extremes of the bronchodilator response (BDR, see below) distribution were selected for whole genome sequencing. Genomic DNA was extracted and purified from whole blood using Wizard^®^ Genomic DNA Purification Kits (Promega, Madison, WI, USA).

### Bronchodilators response measurements

Spirometry was performed and BDR (i.e., ΔFEV_1_, defined as the relative change in FEV_1_) was calculated as previously described [27]. In brief, BDR was calculated according to American Thoracic Society/European Respiratory Society guidelines [82] as the percent change in FEV_1_ after 2 doses of albuterol: that is, BDR = (post-FEV_1_ – pre-FEV_1_) / pre-FEV_1_.

High and low drug responders were selected from the extremes of BDR distribution from GALA II and SAGE II (**Fig 1**). S1 Fig highlighted the BDR distribution of the 1,441 individuals who passed WGS data quality control (see “WGS data processing and quality control”). The ΔFEV_1_ cutoffs for high and low responders are as follows: high responders (ΔFEV_1_ > 16.29 for Puerto Ricans, > 8.55 for Mexicans and > 11.81 for African Americans); low responders (ΔFEV_1_ < 7.23 for Puerto Ricans, < 6.05 for Mexicans and < 5.53 for African Americans).

### Analysis on descriptive data of study subjects

Dichotomous variables were tested for association with BDR using Fisher’s exact test. Continuous variables were tested for normality using the Shapiro-Wilk test. Normally and non-normally distributed continuous variables were tested using Student’s t-test and the Wilcoxon rank-sum test, respectively.

### Sample quality control and whole genome sequencing

DNA samples were quantified by fluorescence using the Quant-iT PicoGreen dsDNA assay (ThermoFisher Scientific, Waltham, MA, USA) on a Spectramax fluorometer (Molecular Devices, Sunnyvale, CA, USA). Sample integrity was ascertained using the Fragment Analyzer™ (Advanced Analytical Technologies, Inc., Ankeny, IA, USA). Samples passing QC were genotyped using the HumanCoreExome-24 array (Illumina^®^, San Diego, CA, USA). Genotyping results were analyzed using VerifyIDintensity [83] to flag sample contamination. Sequencing libraries were constructed using the TruSeq PCR-free DNA HT Library Preparation Kit (Illumina^®^, San Diego, CA, USA) with 500ng DNA input. Briefly, genomic DNA was sheared using a Covaris sonicator (Covaris, Woburn, MA, USA), followed by end-repair and bead-based size selection of fragmented molecules. Selected fragments were then A-tailed and sequence adaptors were ligated onto the fragments, followed by a final bead purification of the libraries. Final libraries were reviewed for size distribution using Fragment Analyzer and quantified by qPCR (Kapa Biosystems, Wilmington, MA, USA). Libraries were sequenced on a HiSeq X system (Illumina^®^, San Diego, CA, USA) with v2 chemistry, using a paired-end read length of 150 bp, to a minimum of 30x mean genome coverage.

### WGS data processing and quality control

Sequencing data were demultiplexed using bcl2fastq version 2.16.0.10 (Illumina^®^, San Diego, CA, USA) and aligned to human reference hs37d5 with decoy sequences using BWA-MEM v0.7.8 [84]. Data were further processed using the GATK best-practices v3.2-2 pipeline [85]. Quality control procedures included marking of duplicate reads using Picard tools v1.83 (http://picard.sourceforge.net), realignment around indels, and base quality recalibration using 1000 Genomes Phase 1 high confidence SNPs, HapMap v3.3, dbSNP v137, 1000 Genomes omni2.5, 1000 Genomes Phase 1 indels, and both Mills and 1000 Genomes gold standard indels. Single-sample genotypes were called using GATK HaplotypeCaller followed by joint genotyping of all subjects. The resulting multi-sample Variant Call Format (VCF) file was used for variant quality score recalibration (VQSR). A 99.8% truth sensitivity tranche level was used for SNPs and 99.0% for indel variants. SNP calls were used to check for sample contamination using VerifyBAMId [83], and sample identity was confirmed by requiring > 99.5% concordance with SNP array (HumanCoreExome-24 array) genotypes.

As part of NIH’s Trans-Omics for Precision Medicine (TOPMed) Program, BAM files were submitted to the Informatics Resource Center (IRC) at the University of Michigan. All 1,484 samples sequenced passed TOPMed’s IRC quality control metrics (mean genome coverage >30X; >95% of genome covered at >10X; and <3% contamination).

VCF-level variants were filtered by GATK version 3.4.46 and VCFtools version 0.1.14 [86]. Variants were filtered according to the following procedures: (1) remove variants that were not indicated as “PASS” in the VCF FILTER column, (2) remove variants in low complexity regions [87] (downloaded from https://github.com/lh3/varcmp/tree/master/scripts/LCR-hs37d5.bed.gz), and (3) keep sample genotypes that have minimum read depths of 10 and genotype qualities of 20 (DP ≥ 10 and GQ ≥ 20). The ratio of homozygous to heterozygous variants (hom/het), ratio of transitions to transversions (Ti/Tv), and other variant summary statistics were generated using GATK VariantEval. VCF files were converted into PLINK format using PLINK 1.9 software [88] according to recommended best practices [89]. Genotype consistency between WGS data and previously published Axiom^®^ Genome-Wide LAT 1 array (Affymetrix, Santa Clara, CA) genotype data (dbGaP phs000920.v1.p1 and phs000921.v1.p1) was assessed using VCFtools [86]. Individuals with percentage consistency three S.D. below the mean (< 96.3%) were removed (N=7, **S10 Fig**). Cryptic relatedness was detected using REAP [90]. Global ancestry and allele frequency used by REAP were estimated using ADMIXTURE in supervised mode [91]. Related individuals (kinship coefficient > 0.044, corresponding to a third degree relationship [92]) were excluded from further analysis (N=36), yielding a final sample size of 1,441 for downstream analysis. Downstream analyses were only performed on biallelic SNPs that passed all quality filters mentioned above and had less than 10% of genotype missingness. The 10% genotype missingness filter was applied per population instead of across all three populations except for the rare variant analysis performed with all three populations combined (see **Methods** section “Multi-variant analyses of combined effects of rare variants on BDR”).

### Principal component analysis

Principal component analysis (PCA) was performed to control for hidden population substructure using EIGENSTRAT’s smartpca program [93]. After using PLINK 1.9 to remove biallelic SNPs with low minor allele frequency (MAF ≤ 0.05) and in linkage disequilibrium (R^2^ > 0.5 in a 50-SNP window with a shift size of 5 SNPs), 710,256 variants remained for input into smartpca.

### Local ancestry estimation

Reference genotypes for European and African ancestries were obtained from the Axiom^®^ Genotype Data Set [94]. SNPs with less than a 95% call rate were removed. Since no Native American reference samples are available in the HapMap database, reference genotypes for Native American ancestry were generated from 71 Native American individuals previously genotyped on the Axiom^®^ Genome-Wide LAT 1 array [27].

To call local ancestry tracts, we first created a subset of our WGS data corresponding to sites found on the Axiom^®^ Genome-Wide LAT 1 array, leaving 765,321 markers. Using PLINK 1.9, we merged these data with our European (CEU), African (YRI), and Native American (NAM) reference panels, which overlapped at 434,145 markers. After filtering multi-allelic SNPs and SNPs with > 10% missing data, we obtained a final merged dataset of 428,644 markers. We phased all samples using SHAPEIT2 [95] and called local ancestry tracts jointly with RFMix [96] under a three-way admixture model based on the African, European, and Native American reference genotypes described above.

### Variant annotation

TOPMed freeze 2 and 3 variants were annotated using the WGSA annotation pipeline [97]. Annotated VCF files were downloaded from the TOPMed Data Coordinating Center SFTP sites. dbSNP150 annotation was added separately by using VCF file downloaded from NCBI dbSNP ftp site [98]. ENCODE (v4) annotations were downloaded as BED files from the UCSC Table Browser (Feb.2009 [GRCh37/hg19] assembly). The conversion from GRCh37 to GRCh38 coordinates was performed using liftOver from the UCSC Genome Browser Utilities [99].

### Single locus BDR association testing on common variants

An additive logistic regression model was used to evaluate the association of biallelic common variants (MAF > 1%) with BDR using PLINK 1.9 separately for each population. Throughout this study, high drug responders were assigned as cases. Logistic regression models included the covariates age, sex and body mass index (BMI) categories to account for previously reported confounders of asthma and BDR [100–107], and the first ten principal components (PCs) to correct for 0 population substructure in admixed populations. BMI and age- and sex-specific BMI percentiles (BMI-pct) were calculated as previously described [49] and used for assignment to BMI categories. For subjects aged 20 years and over, BMI categories were defined as follows: underweight (BMI < 18), normal (18 ≤ BMI < 25), overweight (25 ≤ BMI < 30) and obese (BMI ≥ 30). For subjects under 20 years of age, BMI categories were defined as follows: underweight (BMI-pct < 5), normal (5 ≤ BMI-pct < 85), overweight (85 ≤ BMI-pct < 95) and obese (BMI-pct ≥ 95). Baseline lung function (pre-FEV_1_) has a significant impact on variation in BDR drug response. Pre-FEV_1_ was not included as a covariate in association analyses, as variation in pre-FEV_1_ was indirectly captured by several of the ten principal components already included in association models (S13 Table). A correlation matrix showing the relationship between pre-FEV_1_, age, sex, BMI status, mean global ancestry, and the top ten principal components is presented in S13 Table. The correlation matrix was constructed using Spearman correlation coefficients and the accompanying association tests for the significance of each correlation. Population-specific genome-wide significance thresholds for the single locus analyses were calculated 6based on genotypes using the autocorrelation-based *effectiveSize()* function in the R package *‘coda’* as published by Sobota *et al.* [41]. Population-specific genome-wide significance thresholds after adjusting for the effective number of tests (adjusted genome-wide significance) were 1.57 × 10^-7^ for Puerto Ricans, 2.42 × 10^-7^ for Mexicans, and 9.59 × 10^-8^ for African Americans. Suggestive significance thresholds were calculated as one divided by the effective number of tests [43]. Linkage disequilibrium patterns (Genome build: hg19/1000 Genomes Nov 2014 AMR) of the flanking regions of BDR-associated SNPs were visualized using LocusZoom [108]. Quantile-quantile (q-q) plots were generated using a uniform distribution as the expected *p*-value distribution (**S11 Fig a-c**). The genomic inflation factor (λGC) was calculated using the R package ‘*gap*’.

### Trans-ethnic meta-analysis of common variant effects on BDR

A meta-analysis of the effects of common variants on BDR across the three populations was performed using METASOFT [109]. We used the Han and Eskin random effects model optimized for detecting associations under heterogeneous genetic effects from different study conditions [109]. The number of effective tests was estimated using the R package *‘coda’* as described above, yielding an adjusted genome-wide significance threshold of 3.53 × 10^-7^ and a suggestive significance threshold of 7.06 x 10^-6^. Allele frequency variation in the world population was visualized using the Geography of Genetic Variants Browser (GGV) beta v0.92 [110]. The q-q plot and λ_GC_ were generated in the same way as described above (**S11 Fig d**, see **Methods** section, “Single locus BDR association testing on common variants”).

### Calculation of variation in BDR explained by common variants

Total variation in BDR explained was estimated by calculating McFadden’s pseudo R^2^ [111] in each population separately, after first pruning significantly and suggestively associated variants (**Table 4**) for LD using the LD prune function in PLINK 1.9 (R^2^ cut-off: 0.6, window size = 50 SNPs, shift = 5 SNPs). Calculation of pseudo R^2^ was adjusted for age, sex, BMI category, and principal components 1-10. McFadden’s Pseudo R^2^ is defined as:

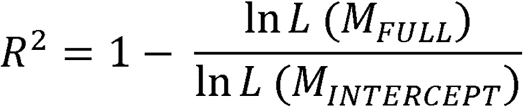

*L*: Estimated likelihood of model

*M_FULL_*: Model with all predictors

*M_INTERCEPT_*: Model with no predictors

### Multi-variant analyses of combined effects of common and rare variants on BDR

Combined effects of common and rare variants on BDR were analyzed using SKAT-O [59]. Common and rare variants were collapsed into 1kb windows sliding across the GRCh37 genome in steps of 500 base pairs. A total of 5.3 million windows were analyzed and the R package *‘coda’* was used to determine the number of effective tests based on autocorrelation of the association *p*-value, as described above. Adjusted genome-wide significance thresholds for Puerto Ricans (8.15 × 10^-8^), Mexicans (8.60 × 10^-8^), African Americans (6.94 × 10^-8^) and for all three populations combined (1.53 × 10^-7^) were used to identify windows of variants significantly associated with BDR. The same covariates used for common variant association testing were used for analyses of individual populations. For analyses of individuals combined across all three populations, we avoided confounding from population substructure by including local ancestry as additional covariates, defined as the proportions of Native American and African ancestries for the window under testing. The q-q plots and λ_GC_ were generated in the same way as described above (**S12 Fig**, see **Methods** section, “Single locus BDR association testing on common variants”).

### Variation in BDR explained by associated SKAT-O regions

Variation in BDR explained by SKAT-O regions was calculated as described above for common variant analyses using McFadden’s pseudo R^2^ [111], but with one addition: variants were weighted using a weighted kernel as described in SKAT-O [112].

### Single locus BDR association and trans-ethnic meta-analysis of array data

To address the discrepancy between our current common variant analysis results with previously published BDR GWAS results [27], we used 1,414 of the 1,441 individuals who had both Axiom^®^ Genome-Wide LAT 1 array (see Methods section, “WGS data processing and QC”) and WGS data available to rerun the single locus BDR association testing and trans-ethnic meta-analysis. Array data were imputed to the Haplotype Reference Consortium [113] (HRC release 1) panel using the Michigan Imputation Server [114]. We used the top 1,000 BDR-associated SNPs to examine the relationship between array-based and WGS-based association *p*-values, genotype discordance, and imputation R^2^. Correlation between the array-based and WGS-based association *p*-values was determined by Spearman correlation. We also performed single locus BDR association testing and trans-ethnic meta-analysis by applying linear regression on 1,122 Puerto Ricans, 662 Mexicans and 1,105 African Americans using BDR (ΔFEV_1_) as a continuous trait. HRC imputed array-based data and the same covariates as described above were used for the analysis.

### Replication of top BDR-associated common variants

Replication cohorts included the Genetics of Asthma in Latino Americans Study (GALA I) [13, 115], the Study of African Americans, Asthma, Genes & Environments (SAGE I) [80], a case-control study of childhood asthma in Puerto Ricans (HPR) [116], the Study of Asthma Phenotypes and Pharmacogenomic Interactions by Race-Ethnicity (SAPPHIRE) [117] and a cohort from the Children’s Hospital of Philadelphia (CHOP) [118]. Descriptive statistics for the replication cohorts are shown in S6 Table.

Logistic regression in replication analyses was performed using the same population-specific extreme BDR cut-offs applied to the discovery analyses in the current study (see Methods section, “Bronchodilator response measurements”). All 27 significantly and suggestively associated SNPs from the discovery analysis (**Table 4**) were assessed in each replication study population. An additive genetic model was assumed for each SNP tested. Association models were adjusted for sex, age, BMI categories, and the first ten principal components as in the discovery analysis. Meta-analysis across all replication studies was performed using METASOFT as described above (Methods, “Trans-ethnic meta-analysis of common variant effects on BDR”).

The GALA I and SAGE I replication cohorts included 108 Puerto Ricans, 202 Mexicans and 141 African Americans with BDR measurements and complete data for all covariates (age, sex, BMI categories and the first ten PCs). Genotype data were imputed to the HRC panel using the Michigan Imputation Server [114]. Replication in the HPR cohort involved 414 Puerto Rican subjects. Spirometry data were collected as previously described [119]. Genome-wide genotyping was performed using the Illumina HumanOmni2.5 BeadChip platform (Illumina Inc., San Diego, CA) and processed as previously described [120]. Genotype data were phased with SHAPE-IT [121] and imputation was performed with IMPUTE2 [122] using all populations from 1000 Genomes Project Phase 3 as reference [75]. The SAPPHIRE replication cohort consisted of 1,022 African Americans with asthma. Genome-wide genotyping was performed using the Axiom^®^ Genome-Wide AFR 1 array (Affymetrix Inc., Santa Clara, CA) as previously described [26]. Genotype data were imputed to the cosmopolitan 1000 Genomes Phase 1 version haplotypes using the Michigan Imputation Server [114]. The CHOP replication cohort included 280 African Americans. Genotyping was performed as described [118], and genotype data were imputed to the HRC panel using the Sanger Imputation server [113].

### Identification of nearest genes for BDR-associated loci

The four nearest transcripts to BDR-associated loci were identified by using the “closest” command in BEDTools with the parameters “-d-k 4” and the RefSeq gene annotations (Feb.2009 [GRCh37/hg19] assembly) downloaded in refFlat format from the UCSC Table Browser [123]. Genes with multiple transcripts were reported only once. When reporting the nearest gene, the “closest” command in BEDTools with the parameters “-D a” was applied.

### Primary bronchial smooth muscle cell culture

Cryopreserved primary human bronchial smooth muscle from two donors (from Lonza catalog number CC-2576, lot number 0000212076 and from ATCC catalog number PCS-130-011, lot number 62326179) was thawed and expanded in Lonza Smooth Muscle Growth Media (SmGM; catalog number CC-3182) on T75 flasks (E&K Scientific Products, catalog number 658175).

### H3K27ac ChIP-seq assay

Upon reaching 80% confluency, BSMCs were serum-starved by replacing SmGM with smooth muscle basal media (SmBM) for 24 hours. BSMCs were then grown in SmBM containing 5% FBS for 4 hours, then fixed in 1% formaldehyde for 10 min and quenched with 0.125 M glycine for 5 minutes. Cells were removed from the T75 flasks by scraping in cold PBS containing sodium butyrate (20 mM, Diagenode, catalog number C12020010). Chromatin sheering was carried out using a Covaris S2 sonicator. Sheared chromatin was used for immunoprecipitation with antibodies against active chromatin marks (H3K27ac; Abcam, ab4729) using the Diagenode LowCell# ChIP kit (CAT#C01010072), following the manufacturer’s protocol. Libraries were prepared using the Rubicon DNA-Seq kit (CAT#R400406) following the manufacturer’s protocol and sequenced on an Illumina HiSeq 4000 using singleend 50-bp reads to a sequencing depth of at least 25 million reads (submitted under BioProject PRJNA369271). Uniquely mapping raw reads were aligned using Bowtie [124] under default settings. Peak regions for each individual were called using MACS2 [125, 126] and reproducible peaks identified using the ENCODE IDR pipeline [127].

### Diverse Convergent Evidence approach for variant prioritization

The Diverse Convergent Evidence (DiCE) approach is a logical, heuristic framework for integrating multiple types of observational, bioinformatics, and laboratory evidence to prioritize variants discovered from high throughput genetic studies for further evaluation in functional experiments [53]. Results from the trans-ethnic meta-analysis and from the replication analyses were considered observational data. Laboratory evidence was provided by the identified peaks in our ChIP-seq analyses performed in BSMCs. Informatic evidence was compiled using Ensembl, PubMed, the NHGRI-EBI GWAS Catalog, and ENCODE to identify previously reported associations with BDR or asthma-related phenotypes, and predicted biological functions associated with assessed loci. After compiling the observational, informatic and laboratory evidence for each suggestively or significantly associated variant, DiCE constructs an evidence matrix to estimate the strength of the information supporting each association. DiCE scores > 6 were considered strong evidence that a given locus was involved in the pathophysiology of BDR, and variants with the highest DiCE score, after meeting this criterion, were prioritized for downstream functional analyses.

### Luciferase assays

*NFKB1* candidate enhancer sequences were amplified from human genomic DNA (Roche) using oligonucleotides designed in Primer3 with 18 and 20 bp overhangs for forward and reverse primers, respectively (5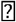-GGCCTAACTGGCCGGTAC-3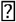 and 5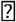-CGCCGAGGCCAGATCTTGAT-3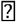), complementary to sequences flanking the KpnI and EcoRV sites in the pGL4.23 Gate A vector (Promega, Madison, WI, USA) using Phusion High-Fidelity PCR kit (NEB, catalog number M0531S). PCR primers were designed around the edges of these ChIP-seq peaks and the most complete fragment that was successfully amplified was used for luciferase assays. PCR products where then cloned into the pGL4.23 vector using the Gibson Assembly method (NEB, catalog number E2611S). Smooth muscle cells were plated at 50-70% confluency in 24-well cell culture plates (Falcon, catalog number 353047) and grown to 80% confluency in SmGM. Transfections were carried out by combining polyethyleneimine (PEI) with DNA vectors at a 1:1 ratio by weight in opti-MEM (Life Technologies, catalog number 31985070). The transfection mixture consisted of 225 ng of enhancer assay vectors and 25 ng of pGL4.24 (Renilla transfection efficiency control) with 250 ng of PEI in 50 μL of opti-MEM. After 15 minutes, 500 μL of SmBM was added to the transfection mixture and the combination added to cell culture. Cells were incubated for 4 hours in SmBM plus the transfection mixture, then media was replaced with SmGM for 24 hours. Cells were then washed with PBS and enhancer assay cells were lysed with 100 μL of Passive Lysis Buffer (Promega, Madison, WI, USA). Reporter activity was measured using the Dual-Luciferase Reporter Assay System (Promega, Madison, WI, USA) and measured on the Glomax 96 well plate luminometer (Promega, Madison, WI, USA). The luciferase assay was carried out in two separate experiments, with three independent replicates per experiment (three wells of cells were transfected per construct per experiment). Each well was then split into two technical replicates for luciferase activity measurements with a luminometer.

### RNA extraction, library preparation and sequencing

Among the African American subjects with WGS data in our study, 39 samples were selected for RNA-seq based on BDR status and the number of copies of low-BDR associated alleles at rs28450894. The number of samples in each category is shown in S14 Table. Peripheral blood samples were collected into PAXgene Blood RNA tubes (PreAnalytiX, Hombrechtikon, Switzerland). Total RNA was extracted from PAXgene Blood RNA tubes using the MagMAX^TM^ for Stabilized Blood Tubes RNA Isolation Kit (CAT#4451894, Thermo Fisher Scientific, Waltham, MA, USA) according to manufacturer’s protocols. RNA integrity and yield were assessed using an Agilent 2100 Bioanalyzer (Agilent Technologies, Santa Clara, CA, USA). Globin depletion was performed using GLOBINclear^TM^ kit (CAT#AM1980, Thermo Fisher Scientific, Waltham, MA, USA). Library preparation and ribosomal depletion were performed using KAPA Stranded RNA-seq Kit with RiboErase (CAT#KK8483, Kapa Biosystems, Wilmington, MA, USA) according to the manufacturer’s protocols. Each sample was uniquely barcoded with NEXTflex^TM^ DNA Barcodes (CAT#514104, Bioo Scientific^®^, Austin, TX, USA). Barcoded libraries were pooled and sequenced on 4 lanes on a HiSeq 4000 sequencing system (Illumina^®^, San Diego, CA, USA) with a paired-end read length of 100 bp at the University of California, San Francisco’s Center for Advanced Technology.

### RNA-seq data processing and analysis

Raw sequencing reads were aligned to the human reference genome (hg19) using STAR (v2.4.2a) [128]. Gene read counts were obtained from uniquely mapped reads based on Ensembl annotation (v75) [129]. DESeq2 [130] was used to analyze read counts for differential gene expression changes between genotypes, including an interaction term with genotype and sex (genotype * sex). We used a linear model to account for sex, age and library prep batch, and a custom model matrix to correct for GC content difference between genes. After normalization for sequencing depth and GC percentage, genes with fewer than an average of five normalized read counts per sample and fewer than 20 samples with at least one read count were removed. This filtering process kept 19,592 Ensembl genes for analysis. Fold change, raw and FDR-adjusted p-value for the genotype term was reported. Genes were then further filtered to analyze the locus surrounding rs28450894 for differential gene expression by including all genes with a transcriptional start site within 1Mbp of rs28450894. *p*-values were then corrected using the false discovery rate method to account for the 13 genes in this locus. Significant level FDR-adjusted *p*-value is ≤ 0.1.

## ACKNOWLEDGEMENTS

Whole genome sequencing (WGS) for the Trans-Omics in Precision Medicine (TOPMed) program was supported by the National Heart, Lung, and Blood Institute (NHLBI). WGS for “NHLBI TOPMed: Genes-environments & Admixture in Latino Americans (GALA II) Study” (phs000920) and “NHLBI TOPMed: Study of African Americans, Asthma, Genes and Environments (SAGE II)” (phs000921) was performed at the New York Genome Center (3R01HL117004-01S3). We acknowledge New York Genome Center investigators and teams for whole genome sequencing sample preparation, quality control, data generation, data processing and initial joint genotyping. Centralized read mapping and genotype calling, along with variant quality metrics and filtering were provided by the TOPMed Informatics Research Center (3R01HL-117626-02S1). Phenotype harmonization, data management, sample-identity QC, and general study coordination were provided by the TOPMed Data Coordinating Center (3R01HL-120393-02S1). We gratefully acknowledge the studies and participants who provided biological samples and data for TOPMed. We also gratefully acknowledge the contributions of the investigators of the NHLBI TOPMed Consortium (https://www.nhlbiwgs.org/topmed-banner-authorship). C.A.W. would like to declare that the content of this publication does not necessarily reflect the views or policies of the Department of Health and Human Services, nor does mention of trade names, commercial products, or organizations imply endorsement by the U.S. Government.

---

**S1 Fig.** Distribution of bronchodilator drug response (BDR) in Puerto Ricans, Mexican (GALA II) and African Americans (SAGE). The 1,441 subjects selected from the extreme of the BDR distribution for this study were highlighted.

**S2 Fig.** Global ancestry composition for **(a)** Puerto Ricans, **(b)** Mexicans and **(c)** African Americans. Each individual is represented by a vertical line and the ancestry composition is colored based on the percentage composition of African (red), European (blue) and Native American (green) ancestries.

**S3 Fig.** Plot of the first two principal components of variation based on WGS genotypes of 1,441 individuals.

**S4 Fig.** Diverse Convergent Evidence (DiCE) prioritization of 27 common variants (Table 4). Y-axis indicates the points allotted per SNP for each form of evidence according to S15 Table: statistical evidence (grey), informatic evidence (blue), and experimental evidence (orange). Evidence-specific scores for each SNP are provided in the table. SNPs with a DiCE score ≥ 4 are labeled with the nearest gene.

**S5 Fig.** Regions overlapped with SNPs in LD with BDR-associated SNPs show enhancer activity in BSMCs. **(a)** H3K27ac ChIP-seq peaks in BSMCs overlap with SNPs in LD with rs28450894 (marked red). The GRCh37 coordinates are chr4:103486504-103491377 for region 1 and chr4:103527184-103531814 for region 2. In addition to BSMCs, H3K27ac data is shown for Roadmap Epigenomic peripheral blood mononuclear cells (PBMSs) and lung tissue, as well as ENCODE project data for GM12878, H1 human embryonic stem cell (H1-ESC) line, human skeletal muscle cells and myoblasts (HSMM) and normal human epidermal keratinocytes (NHEK). **(b)** Luciferase assay results for regions tested for enhancer activity in BSMCs. *NFKB1* region 2 significantly increased the expression of luciferase over empty vector control (Fold change = 2.24, *p* < 0.01). A red line marks a fold change of one compared to the empty vector.

**S6 Fig.** A GGV plot showing the allele frequency of rs2845894 for different populations based on 1000 Genomes Project. European populations include CEU (Utah residents [CEPH] with northern and western ancestry), FIN (Finnish in Finland), GBR (British in England and Scotland), TSI (Toscani in Italia) and IBS (Iberian population in Spain). American populations include MXL (Mexican ancestry from Los Angeles USA), PUR (Puerto Ricans from Puerto Rico), CLM (Colombians from Medellin, Colombia) and PEL (Peruvians from Lima, Peru). African populations include ASW (Americans of African ancestry in SW USA), ACB (African Caribbeans in Barbados), GWD (Gambian in western divisions in the Gambia), MSL (Mende in Sierra Leone), YRI (Yoruba in Ibadan, Nigeria), ESN (Esan in Nigeria) and LWK (Luhya in Webuye, Kenya).

**S7 Fig.** Boxplot showing increasing number of copies of low BDR-associated T allele of rs28450894 is associated with decreased expression of *SLC39A8* in blood regardless of sex (*p* = 0.0066, FDR adjusted *p* = 0.0856, log_2_(β) = -0.327).

**S8 Figure.** Manhattan plot of the single locus BDR association testing for **(a)** Puerto Ricans, **(b)** Mexicans and **(c)** African Americans. The horizontal lines are colored blue for the suggestive significance thresholds and red for the Bonferroni-adjusted genome-wide significance thresholds. Since no associations were close to the Bonferroni-adjusted genome-wide significance thresholds in Mexicans and African Americans, the red line is only marked in Puerto Ricans.

**S9 Fig.** A comparison of the top 1000 BDR associations between the array-based and WGS-based genotype data. **(a)** A plot of association *p*-values of array-based and WGS-based data. The *p*-values from the two data types showed high correlation, especially for SNPs with more significant *p*-values. Genotype discordance of SAGE II **(b)** and GALA II **(c)** WGS SNPs. The corresponding imputation R^2^ of the same SNPs in the HRC imputed array data is indicated by red (R^2^ ≥ 0.8) or blue (R^2^ < 0.8).

**S10 Fig.** Percentage genotype concordance between Axiom LAT1 array and WGS genotypes. Grey horizontal lines mark one, two and three standard deviations (S.D.) from the mean percentage genotype concordance.

**S11 Fig.** Quantile-quantile (q-q) plots of the single locus BDR association for **(a)** Puerto Ricans, **(b)** Mexicans and **(c)** African Americans and **(d)** trans-ethnic metaanalysis. The genomic inflator factors (λ_GC_) are shown on the q-q plots.

**S12 Fig.** Quantile-quantile (q-q) plots of SKAT-O analysis of biallelic common and rare SNPs grouped by 1kb windows for **(a)** Puerto Ricans, **(b)** Mexicans and **(c)** African Americans and **(d)** all individuals in all three populations. The genomic inflator factors (λ_GC_) are shown on the q-q plots.

**S1 Table.** Novel common variants in discovery cohort by population.

**S2 Table.** Chromosomal location of BDR-associated common variants.

**S3 Table.** Common variant BDR association results by population.

**S4 Table.** Lung-related phenotypes previously reported for BDR-associated common variants and their nearest genes.

**S5 Table.** Variation in BDR Status explained by significant and suggestively associated variants identified by trans-ethnic meta-analysis and SKAT-O.

**S6 Table.** Descriptive statistics for replication cohorts.

**S7 Table.** Replication of common variant associations.

**S8 Table.** Size of DNA inserts used in luciferase assay and key variants included within those sequences.

**S9 Table.** Functional annotations for BDR-associated common variants in Table 4.

**S10 Table.** Replication of previously reported BDR-associated SNPs in the current study by population and trans-ethnic meta-analysis.

**S11 Table.** Chromosomal location of identified SKAT-O regions by genome build.

**S12 Table.** Functional annotations for variants within SKAT-O regions.

**S13 Table.** Correlation between baseline lung function, top ten principle components, other covariates included in association analyses in the discovery study population (N =1,441).

**S14 Table.** Number of African American samples selected for RNA-Seq based on BDR status and number of copies of low BDR-associated allele.

**S15 Table.** Diverse Convergent Evidence (DiCE) approach scoring rubric.

## REFERENCES

1. Lara Akinbami. Centers for Disease Control and Prevention. (2015). Asthma Prevalence, Health Care Use and Mortality: United States, 2003-05. [online] Available at: http://www.cdc.gov/nchs/data/hestat/asthma03-05/asthma03-05.htm [Accessed 9/12 2017].

2. Vos T, Flaxman AD, Naghavi M, Lozano R, Michaud C, et al. (2012) Years lived with disability (YLDs) for 1160 sequelae of 289 diseases and injuries 19902010: A systematic analysis for the global burden of disease study 2010. Lancet 380(9859): 2163–2196.

3. World Health Organization. (2017). Asthma. [online] Available at: http://www.who.int/mediacentre/factsheets/fs307/en/ [Accessed 9/12 2017].

4. World Health Organization. (2007). Global surveillance, prevention and control of chronic respiratory diseases: a comprehensive approach. [online] Available at: http://www.who.int/gard/publications/GARD_Manual/en/ [Accessed 9/12 2017].

5. Oh SS, White MJ, Gignoux CR, Burchard EG. (2016) Making precision medicine socially precise. take a deep breath. Am J Respir Crit Care Med 193(4): 348–350.

6. Burchard EG. (2014) Medical research: Missing patients. Nature 513(7518): 301–302.

7. Barr RG, Aviles-Santa L, Davis SM, Aldrich TK, Gonzalez F, II, et al. (2016) Pulmonary disease and age at immigration among hispanics. Results from the Hispanic Community Health Study/Study of Latinos. Am J Respir Crit Care Med 193(4): 386–395.

8. Palmer LJ, Silverman ES, Weiss ST, Drazen JM. (2002) Pharmacogenetics of asthma. Am J Respir Crit Care Med 165(7): 861–866.

9. Nelson HS. (1995) Beta-adrenergic bronchodilators. N Engl J Med 333(8): 499506.

10. Eggleston PA, Malveaux FJ, Butz AM, Huss K, Thompson L, et al. (1998) Medications used by children with asthma living in the inner city. Pediatrics 101(3 Pt 1): 349–354.

11. Finkelstein JA, Lozano P, Farber HJ, Miroshnik I, Lieu TA. (2002) Underuse of controller medications among medicaid-insured children with asthma. Arch Pediatr Adolesc Med 156(6): 562–567.

12. Drazen JM, Silverman EK, Lee TH. (2000) Heterogeneity of therapeutic responses in asthma. Br Med Bull 56(4): 1054–1070.

13. Burchard EG, Avila PC, Nazario S, Casal J, Torres A, et al. (2004) Lower bronchodilator responsiveness in puerto rican than in mexican subjects with asthma. Am J Respir Crit Care Med 169(3): 386–392.

14. Naqvi M, Thyne S, Choudhry S, Tsai HJ, Navarro D, et al. (2007) Ethnic-specific differences in bronchodilator responsiveness among african americans, puerto ricans, and mexicans with asthma. J Asthma 44(8): 639–648.

15. Wechsler ME, Castro M, Lehman E, Chinchilli VM, Sutherland ER, et al. (2011) Impact of race on asthma treatment failures in the asthma clinical research network. Am J Respir Crit Care Med 184(11): 1247–1253.

16. Martinez FD. (2005) Safety of long-acting beta-agonists–an urgent need to clear the air. N Engl J Med 353(25): 2637–2639.

17. Dixon AE. (2011) Long-acting beta-agonists and asthma: The saga continues. Am J Respir Crit Care Med 184(11): 1220–1221.

18. Nelson HS, Weiss ST, Bleecker ER, Yancey SW, Dorinsky PM, et al. (2006) The salmeterol multicenter asthma research trial: A comparison of usual pharmacotherapy for asthma or usual pharmacotherapy plus salmeterol. Chest 129(1): 15–26.

19. Kramer JM. (2009) Balancing the benefits and risks of inhaled long-acting beta-agonists–the influence of values. N Engl J Med 360(16): 1592–1595.

20. McGeachie MJ, Stahl EA, Himes BE, Pendergrass SA, Lima JJ, et al. (2013) Polygenic heritability estimates in pharmacogenetics: Focus on asthma and related phenotypes. Pharmacogenet Genomics 23(6): 324–328.

21. Nieminen MM, Kaprio J, Koskenvuo M. (1991) A population-based study of bronchial asthma in adult twin pairs. Chest 100(1): 70–75.

22. Fagnani C, Annesi-Maesano I, Brescianini S, D’Ippolito C, Medda E, et al. (2008) Heritability and shared genetic effects of asthma and hay fever: An italian study of young twins. Twin Res Hum Genet 11(2): 121–131.

23. Himes BE, Jiang X, Hu R, Wu AC, Lasky-Su JA, et al. (2012) Genome-wide association analysis in asthma subjects identifies SPATS2L as a novel bronchodilator response gene. PLoS Genet 8(7): e1002824.

24. Duan QL, Lasky-Su J, Himes BE, Qiu W, Litonjua AA, et al. (2014) A genomewide association study of bronchodilator response in asthmatics. Pharmacogenomics J 14(1): 41–47.

25. Israel E, Lasky-Su J, Markezich A, Damask A, Szefler SJ, et al. (2015) Genomewide association study of short-acting beta2-agonists. A novel genome-wide significant locus on chromosome 2 near ASB3. Am J Respir Crit Care Med 191(5): 530–537.

26. Padhukasahasram B, Yang JJ, Levin AM, Yang M, Burchard EG, et al. (2014) Gene-based association identifies SPATA13-AS1 as a pharmacogenomic predictor of inhaled short-acting beta-agonist response in multiple population groups. Pharmacogenomics J 14(4): 365–371.

27. Drake KA, Torgerson DG, Gignoux CR, Galanter JM, Roth LA, et al. (2014) A genome-wide association study of bronchodilator response in latinos implicates rare variants. J Allergy Clin Immunol 133(2): 370–378.

28. Conrad DF, Jakobsson M, Coop G, Wen X, Wall JD, et al. (2006) A worldwide survey of haplotype variation and linkage disequilibrium in the human genome. Nat Genet 38(11): 1251–1260.

29. Hoffmann TJ, Zhan Y, Kvale MN, Hesselson SE, Gollub J, et al. (2011) Design and coverage of high throughput genotyping arrays optimized for individuals of east asian, african american, and latino race/ethnicity using imputation and a novel hybrid SNP selection algorithm. Genomics 98(6): 422–430.

30. Illumina. (2016). Infinium^®^ Multi-Ethnic Global BeadChip. [online] Available at: https://www.illumina.com/content/dam/illumina-marketing/documents/products/datasheets/multi-ethnic-global-data-sheet-370-2016-001.pdf [Accessed 9/12 2017].

31. Zheng HF, Rong JJ, Liu M, Han F, Zhang XW, et al. (2015) Performance of genotype imputation for low frequency and rare variants from the 1000 genomes. PLoS One 10(1): e0116487.

32. Huang J, Howie B, McCarthy S, Memari Y, Walter K, et al. (2015) Improved imputation of low-frequency and rare variants using the UK10K haplotype reference panel. Nat Commun 6: 8111.

33. Zhang F, Lupski JR. (2015) Non-coding genetic variants in human disease. Hum Mol Genet 24(R1): R102–10.

34. Bustamante CD, Burchard EG, De la Vega FM. (2011) Genomics for the world. Nature 475(7355): 163–165.

35. Popejoy AB, Fullerton SM. (2016) Genomics is failing on diversity. Nature 538(7624): 161–164.

36. Oh SS, Galanter J, Thakur N, Pino-Yanes M, Barcelo NE, et al. (2015) Diversity in clinical and biomedical research: A promise yet to be fulfilled. PLoS Med 12(12): e1001918.

37. Hankinson JL, Odencrantz JR, Fedan KB. (1999) Spirometric reference values from a sample of the general U.S. population. Am J Respir Crit Care Med 159(1): 179–187.

38. Kircher M, Witten DM, Jain P, O’Roak BJ, Cooper GM, et al. (2014) A general framework for estimating the relative pathogenicity of human genetic variants. Nat Genet 46(3): 310–315.

39. Gulko B, Hubisz MJ, Gronau I, Siepel A. (2015) A method for calculating probabilities of fitness consequences for point mutations across the human genome. Nat Genet 47(3): 276–283.

40. Altshuler D, Daly MJ, Lander ES. (2008) Genetic mapping in human disease. Science 322(5903): 881–888.

41. Sobota RS, Shriner D, Kodaman N, Goodloe R, Zheng W, et al. (2015) Addressing population-specific multiple testing burdens in genetic association studies. Ann Hum Genet 79(2): 136–147.

42. Pe’er I, Yelensky R, Altshuler D, Daly MJ. (2008) Estimation of the multiple testing burden for genomewide association studies of nearly all common variants. Genet Epidemiol 32(4): 381–385.

43. Duggal P, Gillanders EM, Holmes TN, Bailey-Wilson JE. (2008) Establishing an adjusted *p*-value threshold to control the family-wide type 1 error in genome wide association studies. BMC Genomics 9: 516–2164-9-516.

44. Wang H, Thomas DC, Pe’er I, Stram DO. (2006) Optimal two-stage genotyping designs for genome-wide association scans. Genet Epidemiol 30(4): 356–368.

45. Skol AD, Scott LJ, Abecasis GR, Boehnke M. (2007) Optimal designs for two-stage genome-wide association studies. Genet Epidemiol 31(7): 776–788.

46. Reed E, Nunez S, Kulp D, Qian J, Reilly MP, et al. (2015) A guide to genomewide association analysis and post-analytic interrogation. Stat Med 34(28): 3769–3792.

47. Zaykin DV, Zhivotovsky LA. (2005) Ranks of genuine associations in whole-genome scans. Genetics 171(2): 813–823.

48. Castro-Giner F, de Cid R, Gonzalez JR, Jarvis D, Heinrich J, et al. (2010) Positionally cloned genes and age-specific effects in asthma and atopy: An international population-based cohort study (ECRHS). Thorax 65(2): 124–131.

49. White MJ, Risse-Adams O, Goddard P, Contreras MG, Adams J, et al. (2016) Novel genetic risk factors for asthma in african american children: Precision medicine and the SAGE II study. Immunogenetics 68(6-7): 391–400.

50. Johnson M. (1998) The beta-adrenoceptor. Am J Respir Crit Care Med 158(5 Pt 3): S146–53.

51. Nuzzo R. (2014) Scientific method: Statistical errors. Nature 506(7487): 150152.

52. Malley JD, Dasgupta A, Moore JH. (2013) The limits of *p*-values for biological data mining. BioData Min 6(1): 10-0381-6-10.

53. Ciesielski TH, Pendergrass SA, White MJ, Kodaman N, Sobota RS, et al. (2014) Diverse convergent evidence in the genetic analysis of complex disease: Coordinating omic, informatic, and experimental evidence to better identify and validate risk factors. BioData Min 7: 10-0381-7-10. eCollection 20142.

54. Cumberland PM, Czanner G, Bunce C, Dore CJ, Freemantle N, et al. (2014) Ophthalmic statistics note: The perils of dichotomising continuous variables. Br J Ophthalmol 98(6): 841–843.

55. Peloso GM, Rader DJ, Gabriel S, Kathiresan S, Daly MJ, et al. (2016) Phenotypic extremes in rare variant study designs. Eur J Hum Genet 24(6): 924–930.

56. Li D, Lewinger JP, Gauderman WJ, Murcray CE, Conti D. (2011) Using extreme phenotype sampling to identify the rare causal variants of quantitative traits in association studies. Genet Epidemiol 35(8): 790–799.

57. Guey LT, Kravic J, Melander O, Burtt NP, Laramie JM, et al. (2011) Power in the phenotypic extremes: A simulation study of power in discovery and replication of rare variants. Genet Epidemiol 35(4): 236–246.

58. Lamina C. (2011) Digging into the extremes: A useful approach for the analysis of rare variants with continuous traits? BMC Proc 5 Suppl 9: S105-6561-5-S9-S105.

59. Lee S, Wu MC, Lin X. (2012) Optimal tests for rare variant effects in sequencing association studies. Biostatistics 13(4): 762–775.

60. Lee JH, McDonald ML, Cho MH, Wan ES, Castaldi PJ, et al. (2014) *DNAH5* is associated with total lung capacity in chronic obstructive pulmonary disease. Respir Res 15: 97-014-0097-y.

61. Ortiz RA, Barnes KC. (2015) Genetics of allergic diseases. Immunol Allergy Clin North Am 35(1): 19–44.

62. Ramasamy A, Curjuric I, Coin LJ, Kumar A, McArdle WL, et al. (2011) A genome-wide meta-analysis of genetic variants associated with allergic rhinitis and grass sensitization and their interaction with birth order. J Allergy Clin Immunol 128(5): 996–1005.

63. Naqvi M, Choudhry S, Tsai HJ, Thyne S, Navarro D, et al. (2007) Association between IgE levels and asthma severity among african american, mexican, and puerto rican patients with asthma. J Allergy Clin Immunol 120(1): 137–143.

64. Edwards MR, Bartlett NW, Clarke D, Birrell M, Belvisi M, et al. (2009) Targeting the NF-kappaB pathway in asthma and chronic obstructive pulmonary disease. Pharmacol Ther 121(1): 1–13.

65. Zhen G, Park SW, Nguyenvu LT, Rodriguez MW, Barbeau R, et al. (2007) IL-13 and epidermal growth factor receptor have critical but distinct roles in epithelial cell mucin production. Am J Respir Cell Mol Biol 36(2): 244–253.

66. Murgia C, Grosser D, Truong-Tran AQ, Roscioli E, Michalczyk A, et al. (2011) Apical localization of zinc transporter ZnT4 in human airway epithelial cells and its loss in a murine model of allergic airway inflammation. Nutrients 3(11): 910928.

67. Besecker B, Bao S, Bohacova B, Papp A, Sadee W, et al. (2008) The human zinc transporter SLC39A8 (Zip8) is critical in zinc-mediated cytoprotection in lung epithelia. Am J Physiol Lung Cell Mol Physiol 294(6): L1127–36.

68. Persson H, Kwon AT, Ramilowski JA, Silberberg G, Soderhall C, et al. (2015) Transcriptome analysis of controlled and therapy-resistant childhood asthma reveals distinct gene expression profiles. J Allergy Clin Immunol 136(3): 638648.

69. Lin YJ, Chang JS, Liu X, Tsang H, Chien WK, et al. (2015) Genetic variants in PLCB4/PLCB1 as susceptibility loci for coronary artery aneurysm formation in kawasaki disease in han chinese in taiwan. Sci Rep 5: 14762.

70. Hersch M, Peter B, Kang HM, Schupfer F, Abriel H, et al. (2012) Mapping genetic variants associated with beta-adrenergic responses in inbred mice. PLoS One 7(7): e41032.

71. Dougherty RH, Sidhu SS, Raman K, Solon M, Solberg OD, et al. (2010) Accumulation of intraepithelial mast cells with a unique protease phenotype in T(H)2-high asthma. J Allergy Clin Immunol 125(5): 1046–1053.e8.

72. Woodruff PG, Boushey HA, Dolganov GM, Barker CS, Yang YH, et al. (2007) Genome-wide profiling identifies epithelial cell genes associated with asthma and with treatment response to corticosteroids. Proc Natl Acad Sci U S A 104(40): 15858–15863.

73. Motsinger-Reif AA, Jorgenson E, Relling MV, Kroetz DL, Weinshilboum R, et al. (2013) Genome-wide association studies in pharmacogenomics: Successes and lessons. Pharmacogenet Genomics 23(8): 383–394.

74. Tishkoff SA, Reed FA, Friedlaender FR, Ehret C, Ranciaro A, et al. (2009) The genetic structure and history of africans and african americans. Science 324(5930): 1035–1044.

75. 1000 Genomes Project Consortium, Auton A, Brooks LD, Durbin RM, Garrison EP, et al. (2015) A global reference for human genetic variation. Nature 526(7571): 68–74.

76. Zou JY, Park DS, Burchard EG, Torgerson DG, Pino-Yanes M, et al. (2015) Genetic and socioeconomic study of mate choice in latinos reveals novel assortment patterns. Proc Natl Acad Sci U S A 112(44): 13621–13626.

77. Hall SS. (2013) Genetics: A gene of rare effect. Nature 496(7444): 152–155.

78. Burchard EG, Oh SS, Foreman MG, Celedon JC. (2015) Moving toward true inclusion of racial/ethnic minorities in federally funded studies. A key step for achieving respiratory health equality in the united states. Am J Respir Crit Care Med 191(5): 514–521.

79. Borrell LN, Nguyen EA, Roth LA, Oh SS, Tcheurekdjian H, et al. (2013) Childhood obesity and asthma control in the GALA II and SAGE II studies. Am J Respir Crit Care Med 187(7): 697–702.

80. Nishimura KK, Galanter JM, Roth LA, Oh SS, Thakur N, et al. (2013) Early-life air pollution and asthma risk in minority children. the GALA II and SAGE II studies. Am J Respir Crit Care Med 188(3): 309–318.

81. Thakur N, Oh SS, Nguyen EA, Martin M, Roth LA, et al. (2013) Socioeconomic status and childhood asthma in urban minority youths. the GALA II and SAGE II studies. Am J Respir Crit Care Med 188(10): 1202–1209.

82. Pellegrino R, Viegi G, Brusasco V, Crapo RO, Burgos F, et al. (2005) Interpretative strategies for lung function tests. Eur Respir J 26(5): 948–968.

83. Jun G, Flickinger M, Hetrick KN, Romm JM, Doheny KF, et al. (2012) Detecting and estimating contamination of human DNA samples in sequencing and array-based genotype data. Am J Hum Genet 91(5): 839–848.

84. Li H, Durbin R. (2009) Fast and accurate short read alignment with burrowswheeler transform. Bioinformatics 25(14): 1754–1760.

85. DePristo MA, Banks E, Poplin R, Garimella KV, Maguire JR, et al. (2011) A framework for variation discovery and genotyping using next-generation DNA sequencing data. Nat Genet 43(5): 491–498.

86. Danecek P, Auton A, Abecasis G, Albers CA, Banks E, et al. (2011) The variant call format and VCFtools. Bioinformatics 27(15): 2156–2158.

87. Li H. (2014) Toward better understanding of artifacts in variant calling from high-coverage samples. Bioinformatics 30(20): 2843–2851.

88. Chang CC, Chow CC, Tellier LC, Vattikuti S, Purcell SM, et al. (2015) Second-generation PLINK: Rising to the challenge of larger and richer datasets. Gigascience 4: 7-015-0047-8. eCollection 2015.

89. Genetics For Fun. Best practice for converting VCF files to plink format. [online] Available at: http://apol1.blogspot.nl/2014/11/best-practice-for-converting-vcf-files.html.

90. Thornton T, Tang H, Hoffmann TJ, Ochs-Balcom HM, Caan BJ, et al. (2012) Estimating kinship in admixed populations. Am J Hum Genet 91(1): 122–138.

91. Alexander DH, Novembre J, Lange K. (2009) Fast model-based estimation of ancestry in unrelated individuals. Genome Res 19(9): 1655–1664.

92. Manichaikul A, Mychaleckyj JC, Rich SS, Daly K, Sale M, et al. (2010) Robust relationship inference in genome-wide association studies. Bioinformatics 26(22): 2867–2873.

93. Price AL, Patterson NJ, Plenge RM, Weinblatt ME, Shadick NA, et al. (2006) Principal components analysis corrects for stratification in genome-wide association studies. Nat Genet 38(8): 904–909.

94. Affymetrix. (2014). Axiom genotype data set. [online] Available at: http://www.affymetrix.com/support/technical/sample_data/axiom_db/axiomdb_data.affx [Accessed 4/1 2014].

95. Delaneau O, Zagury JF, Marchini J. (2013) Improved whole-chromosome phasing for disease and population genetic studies. Nat Methods 10(1): 5–6.

96. Maples BK, Gravel S, Kenny EE, Bustamante CD. (2013) RFMix: A discriminative modeling approach for rapid and robust local-ancestry inference. Am J Hum Genet 93(2): 278–288.

97. Liu X, White S, Peng B, Johnson AD, Brody JA, et al. (2016) WGSA: An annotation pipeline for human genome sequencing studies. J Med Genet 53(2): 111–112.

98. Sherry ST, Ward MH, Kholodov M, Baker J, Phan L, et al. (2001) dbSNP: The NCBI database of genetic variation. Nucleic Acids Res 29(1): 308–311.

99. Kuhn RM, Haussler D, Kent WJ. (2013) The UCSC genome browser and associated tools. Brief Bioinform 14(2): 144–161.

100. Yao TC, Ou LS, Yeh KW, Lee WI, Chen LC, et al. (2011) Associations of age, gender, and BMI with prevalence of allergic diseases in children: PATCH study. J Asthma 48(5): 503–510.

101. Nicolai T, Pereszlenyiova-Bliznakova L, Illi S, Reinhardt D, von Mutius E. (2003) Longitudinal follow-up of the changing gender ratio in asthma from childhood to adulthood: Role of delayed manifestation in girls. Pediatr Allergy Immunol 14(4): 280–283.

102. Joseph M, Elliott M, Zelicoff A, Qian Z, Trevathan E, et al. (2016) Racial disparity in the association between body mass index and self-reported asthma in children: A population-based study. J Asthma 53(5): 492–497.

103. Dixon AE, Shade DM, Cohen RI, Skloot GS, Holbrook JT, et al. (2006) Effect of obesity on clinical presentation and response to treatment in asthma. J Asthma 43(7): 553–558.

104. Ullah MI, Newman GB, Saunders KB. (1981) Influence of age on response to ipratropium and salbutamol in asthma. Thorax 36(7): 523–529.

105. Mohamed MH, Lima JJ, Eberle LV, Self TH, Johnson JA. (1999) Effects of gender and race on albuterol pharmacokinetics. Pharmacotherapy 19(2): 157161.

106. Carroll CL, Bhandari A, Zucker AR, Schramm CM. (2006) Childhood obesity increases duration of therapy during severe asthma exacerbations. Pediatr Crit Care Med 7(6): 527–531.

107. Forno E, Lescher R, Strunk R, Weiss S, Fuhlbrigge A, et al. (2011) Decreased response to inhaled steroids in overweight and obese asthmatic children. J Allergy Clin Immunol 127(3): 741–749.

108. Pruim RJ, Welch RP, Sanna S, Teslovich TM, Chines PS, et al. (2010) LocusZoom: Regional visualization of genome-wide association scan results. Bioinformatics 26(18): 2336–2337.

109. Han B, Eskin E. (2011) Random-effects model aimed at discovering associations in meta-analysis of genome-wide association studies. Am J Hum Genet 88(5): 586–598.

110. Marcus JH, Novembre J. (2017) Visualizing the geography of genetic variants. Bioinformatics 33(4): 594–595.

111. McFadden D. (1979) Quantitative methods for analysing travel behavior of individuals: Some recent developments. In: Hensher DA, Stopher PR, editors. Behavioural travel modelling. London: Croom Helm. pp. 279–318.

112. Lee S, Emond MJ, Bamshad MJ, Barnes KC, Rieder MJ, et al. (2012) Optimal unified approach for rare-variant association testing with application to small-sample case-control whole-exome sequencing studies. Am J Hum Genet 91(2): 224–237.

113. McCarthy S, Das S, Kretzschmar W, Delaneau O, Wood AR, et al. (2016) A reference panel of 64,976 haplotypes for genotype imputation. Nat Genet 48(10): 1279–1283.

114. Das S, Forer L, Schonherr S, Sidore C, Locke AE, et al. (2016) Next-generation genotype imputation service and methods. Nat Genet 48(10): 1284–1287.

115. Torgerson DG, Gignoux CR, Galanter JM, Drake KA, Roth LA, et al. (2012) Case-control admixture mapping in latino populations enriches for known asthma-associated genes. J Allergy Clin Immunol 130(1): 76–82.e12.

116. Han YY, Forno E, Brehm JM, Acosta-Perez E, Alvarez M, et al. (2015) Diet, interleukin-17, and childhood asthma in puerto ricans. Ann Allergy Asthma Immunol 115(4): 288–293.e1.

117. Levin AM, Wang Y, Wells KE, Padhukasahasram B, Yang JJ, et al. (2014) Nocturnal asthma and the importance of race/ethnicity and genetic ancestry. Am J Respir Crit Care Med 190(3): 266–273.

118. Ong BA, Li J, McDonough JM, Wei Z, Kim C, et al. (2013) Gene network analysis in a pediatric cohort identifies novel lung function genes. PLoS One 8(9): e72899.

119. Miller MR, Hankinson J, Brusasco V, Burgos F, Casaburi R, et al. (2010) Standardisation of lung function testing: The authors’ replies to readers’ comments. Eur Respir J 36(6): 1496–1498.

120. Chen W, Brehm JM, Lin J, Wang T, Forno E, et al. (2015) Expression quantitative trait loci (eQTL) mapping in puerto rican children. PLoS One 10(3): e0122464.

121. Delaneau O, Marchini J, Zagury JF. (2011) A linear complexity phasing method for thousands of genomes. Nat Methods 9(2): 179–181.

122. Howie BN, Donnelly P, Marchini J. (2009) A flexible and accurate genotype imputation method for the next generation of genome-wide association studies. PLoS Genet 5(6): e1000529.

123. Karolchik D, Hinrichs AS, Furey TS, Roskin KM, Sugnet CW, et al. (2004) The UCSC table browser data retrieval tool. Nucleic Acids Res 32 (Database issue): D493–6.

124. Langmead B, Trapnell C, Pop M, Salzberg SL. (2009) Ultrafast and memory-efficient alignment of short DNA sequences to the human genome. Genome Biol 10(3): R25-2009-10-3-r25. Epub 2009 Mar 4.

125. Zhang Y, Liu T, Meyer CA, Eeckhoute J, Johnson DS, et al. (2008) Model-based analysis of ChIP-seq (MACS). Genome Biol 9(9): R137-2008-9-9-r137. Epub 2008 Sep 17.

126. Feng J, Liu T, Qin B, Zhang Y, Liu XS. (2012) Identifying ChIP-seq enrichment using MACS. Nat Protoc 7(9): 1728–1740.

127. Landt SG, Marinov GK, Kundaje A, Kheradpour P, Pauli F, et al. (2012) ChIP-seq guidelines and practices of the ENCODE and modENCODE consortia. Genome Res 22(9): 1813–1831.

128. Dobin A, Davis CA, Schlesinger F, Drenkow J, Zaleski C, et al. (2013) STAR: Ultrafast universal RNA-seq aligner. Bioinformatics 29(1): 15–21.

129. Flicek P, Amode MR, Barrell D, Beal K, Billis K, et al. (2014) Ensembl 2014. Nucleic Acids Res 42(Database issue): D749–55.

130. Love MI, Huber W, Anders S. (2014) Moderated estimation of fold change and dispersion for RNA-seq data with DESeq2. Genome Biol 15(12): 550.

